# No effects of APOE ε4 on spatial navigation and broader cognition in young adults genetically at risk for Alzheimer’s disease

**DOI:** 10.64898/2026.07.10.737269

**Authors:** Luise P. Graichen, Lea Schenk, Christian Gausterer, Isabella C. Wagner

## Abstract

Alzheimer’s disease (AD) causes progressive memory loss and disorientation. It is preceded by a prolonged preclinical phase marked by pathological changes in medial temporal lobe regions involved in spatial navigation. Spatial navigation tasks have been proposed for early AD detection, and altered navigation performance was reported in older carriers of the apolipoprotein E (APOE) ε4 allele, the major genetic risk factor for sporadic AD. However, whether spatial navigation or other cognitive abilities are affected in younger ε4 carriers remains unclear. Here, we genotyped 1000 healthy young adults (18-35 years) who completed the app-based navigation game “Sea Hero Quest” and several tasks assessing working memory, processing speed, executive functioning, and face recognition. ε4 carriers (ε3ε4, N = 88) showed no significant differences from non-carriers (ε3ε3, N = 327) in spatial navigation or other cognitive abilities, supported by equivalence testing and Bayesian analyses. Exploratory findings suggested altered spatial navigation in ε2 carriers (ε2ε2/ε2ε3/ε2ε4, Ns = 7/51/7) versus ε3ε3 controls, who stayed closer to environmental borders and showed better memory updating, face recognition, and processing speed. Therefore, APOE-related behavioural differences in young adults appear small at best, highlighting the need for paradigms sensitive to very subtle changes decades before potential dementia onset.

## Introduction

Alzheimer’s disease (AD) is the most common type of dementia, characterised by progressive memory loss and disorientation. With a rapidly ageing population, its prevalence continues to rise, placing a growing economic and societal burden on healthcare systems worldwide [1,2]. Although clinical symptoms typically emerge around age 65, the underlying pathological changes, such as the accumulation of amyloid beta plaques and tau neurofibrillary tangles in hippocampal and entorhinal regions (critical for memory and spatial navigation) can precede overt disease onset by several decades [3,4]. There is currently no cure for AD, but available treatments aim at slowing disease progression once symptoms appear. Most individuals, however, remain undiagnosed throughout the preclinical stage during which intervention could be most effective [5].

The apolipoprotein E (APOE) gene is the major genetic risk factor for sporadic AD and occurs in three allelic variants (ε2, ε3, and ε4) [6]. AD risk scales with the number of ε4 alleles, increasing approximately 2-3-fold with one copy and 9-12 fold with two [6,7] (but note that these estimates vary with age, sex, and ethnicity [8]). Cognitive-behavioural tasks taxing spatial navigation have been shown to detect ε4-related changes, potentially offering tools for early AD detection [9–13]. For instance, Coughlan et al. [14] used the app-based game “Sea Hero Quest” (SHQ) to assess spatial navigation in middle-aged and older adults at genetic risk for AD (50-70 years), evaluating their performance against benchmark data from more than 27,000 players (50-75 years). They found that ε4 carriers were poorer navigators than ε3 controls (who are not at increased AD risk). Bierbrauer et al. [15] tested navigation ability using a desktop-based homing task and reported impaired path integration in ε4 carriers (18-75 years), suggesting a deficit in tracking one’s position based on self-motion cues [16]. This deficit was more pronounced in older individuals (42-75 years), but appeared largely unimpaired in younger participants (18-41 years). Importantly, Kunz et al. [17] investigated younger ε4 carriers (18-31 years) and reported reduced grid-like codes in the entorhinal cortex, known to subserve path integration [18], despite no behavioural navigation impairment. Together, these findings suggest that spatial navigation tasks may capture ε4-related changes before clinical symptoms emerge, but findings in younger adults are less clear and might be hampered by small sample sizes. Building on the approach by Coughlan et al. [14], we used the SHQ game to examine spatial navigation in a large-scale sample of healthy young ε4 carriers, reasoning that any behavioural deficits in these at-risk individuals could aid future interventions aimed at delaying disease onset from an early age.

Beyond spatial navigation, AD symptoms affect a range of other cognitive domains, including working memory, executive functioning, and face recognition [19–21]. Deficits in working memory and executive functioning have been detected in older individuals (60-90 years) with preclinical AD [22]. During later disease stages, symptoms include impaired face recognition, with some patients (63-87 years) no longer recognising close family members [23]. Evidence for APOE genotype effects across these domains outside clinical populations, however, remains mixed and often contradictory [24–28]. Older ε4 carriers have shown poorer executive functioning, such as attentional shifting (41-85 years) [7] and working memory deficits (39-79 years) [29]. Findings in younger ε4 carriers remain mixed, with some studies reporting no performance differences in samples aged 9-31 years [25] or 35-60 years [30] and others reporting cognitive benefits. For instance, young ε4 carriers (18-32 years) outperformed non-carriers on a choice reaction task as executive control demands increased [31]. This aligns with an account of antagonistic pleiotropy, which proposes early-life cognitive benefits of the ε4 allele that become detrimental later in life. We therefore probed working memory, executive functioning, and face recognition in the same large-scale sample to clarify the role of APOE ε4 for broader cognition.

In the present study, we genotyped 1,000 healthy young adults and identified ε4 carriers (ε3ε4, N = 173, 17.3%) and non-carriers (ε3ε3, N = 669, 66.9%). Participants completed the app-based game Sea Hero Quest (SHQ) on their smartphones, along with cognitive tasks and questionnaires assessing working memory (memory updating, MU; spatial short-term memory, SSTM), executive function (Trail Making Test; TMT A & B), and face recognition (Cambridge Face Memory Test; CFMT). Based on previous findings [14], we hypothesised that healthy young ε3ε4 carriers would show reduced spatial navigation performance relative to ε3ε3 controls. We further examined whether ε4 status was associated with altered working memory, executive function, and face recognition. Lastly, while APOE ε4 has been widely studied as a risk allele for AD, substantially less is known about APOE ε2. In older adults, APOE ε2 was associated with lower AD risk, delayed disease onset, and reduced AD-related neuropathology [32–34]. We therefore extended our analysis to examine whether APOE ε2 (ε2ε2, N = 18; ε2ε3, N = 105; ε2ε4, N = 13) was associated with better spatial navigation and cognitive abilities in healthy young adults compared to controls.

## Results

### Study sample and APOE genotype groups

From an initial sample of 1,000 participants, we excluded 585 individuals due to incomplete data, self-reported neurological or psychiatric conditions, or failure to carry the APOE genotypes of interest (see Figure 1 and Methods). This yielded a final sample of N = 415 participants for the SHQ analyses at the main focus of the present study (327 APOE ε3ε3 and ε3ε4 carriers, consistent with genotype prevalences reported in the general population [35]). Genotype groups showed no significant differences in age, biological sex, or education (all p > .05; Table 1). For the remaining cognitive tasks assessing working memory, processing speed, executive function, and face recognition, sample sizes varied slightly between tasks since not all participants completed all study components (ε3ε3/ε3ε4, Memory Updating Task: n = 308/84, Spatial Short-Term Memory Task: n = 306/83, Cambridge Face Memory Test: n = 318/88, Trail Making Test A & B: n = 335/92). An overview of all genotype groups and exclusion steps is provided in Figure 1.

**Figure 1.**
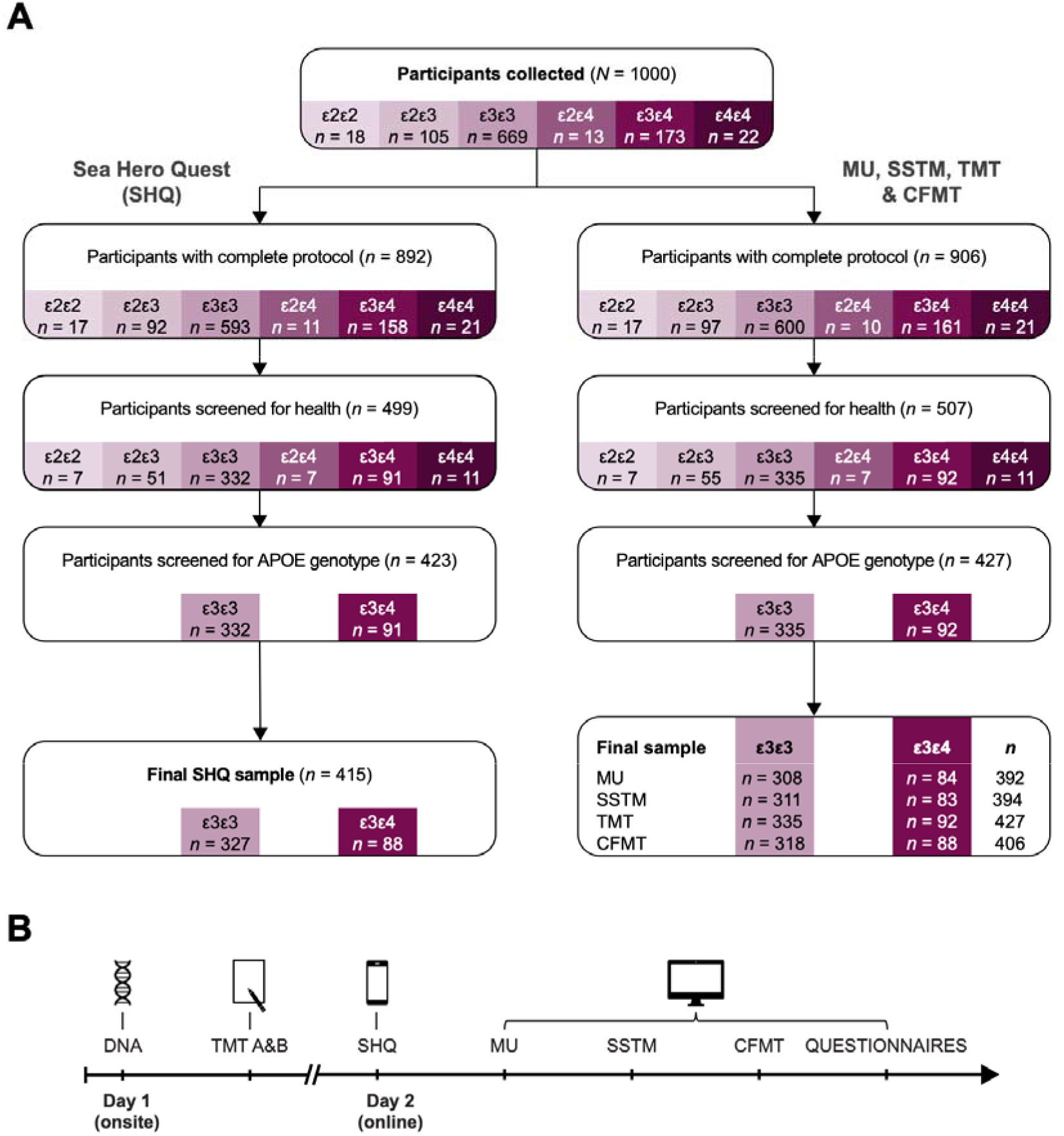
Study procedure and sample selection. A. Sample selection for the SHQ (left) and the other cognitive tasks (right). The final samples were derived in four steps. First, of the 1,000 participants who enrolled, those who did not complete all study components (e.g., the questionnaires) were excluded. Second, participants reporting a psychiatric or neurological history were excluded. Third, for the main analyses, only participants carrying one of the required genotypes (APOE ε3ε3 or ε3ε4) were retained. Fourth, participants were excluded if they did not complete all SHQ levels (SHQ branch) or did not provide valid task responses (cognitive task branch). The diagram shows the number of participants per genotype remaining after each exclusion step. B. Overview of the study procedure. On Day 1 (onsite), participants provided a DNA sample via buccal swab an completed the TMT A & B. On Day 2 (online), they completed the app-based navigation game SHQ, three additional tasks assessing working memory (MU, SSTM) and face recognition (CFMT), and a set of questionnaires.

**Table 1.** Demographics of the main APOE genotype groups.

| | | APOE $\epsilon 3\epsilon 3$ carriers ($n = 327$) | APOE $\epsilon 3\epsilon 4$ carriers ($n = 88$) |
| --- | --- | --- | --- |
| Age | Mean (SD) | 23.21 (3.58) | 23.47 (3.62) |
| Biological sex | Female | 200 (61.2%) | 62 (70.5%) |
|  | Male | 127 (38.8%) | 26 (29.5%) |
| Education | High school | 37 (11.3%) | 11 (12.5%) |
|  | College | 133 (41.0%) | 36 (40.9%) |
|  | University | 150 (45.9%) | 41 (46.6%) |

*Note.* These participants completed the SHQ game. Group differences were tested using a Welch two-sample $t$ -test for age and Pearson's $\chi^2$ tests for sex and education.
Age: $t(136.51) = -0.59$ , $p = .557$ , Cohen's $d = -0.07$ , 95% CI [-0.31, 0.17].
Sex: $\chi^2(1) = 2.19$ , $p = .139$ , adjusted Cramer's $v = 0.06$ , 95% CI [0.00, 1.00].
Education (highest qualification obtained according to German education levels, with High school = Realschulabschluss, College = Gymnasialabschluss): $\chi^2(2) = 0.06$ , $p = .971$ , adjusted Cramer's $v = 0.00$ , 95% CI [0.00, 1.00].

### No significant effects of APOE genotype on wayfinding distance, wayfinding duration, and mean distance-to-border in the SHQ game

We next turned to our main analysis and examined whether the APOE genotype was associated with spatial navigation performance in the SHQ game. Spatial navigation relies on medial temporal lobe regions affected early in AD and has been proposed as a sensitive behavioural marker of early AD- and APOE ε4-related cognitive change [3,4,11]. While differences have been reported between middle-aged APOE ε4 carriers and non-carriers [14,15], spatial navigation differences in younger ε4 carriers are less clear [17]. We therefore examined whether SHQ performance would differ between healthy young ε4 carriers and non-carriers.

While playing the SHQ game, participants navigated a virtual boat through different river landscapes (reflecting different levels of difficulty). At the beginning of each level, they viewed a map displaying several checkpoints (i.e., goals indicated by buoys with flags; allocentric view; Figure 2A, left panel) and were asked to reach them in a specific order. The game then switched to an egocentric view, instructing participants to navigate to the same checkpoints (Figure 2A, middle panel). Spatial navigation performance was quantified using two primary wayfinding outcomes: the distance travelled to complete a level (wayfinding distance; skilled navigators should take shorter routes) and the time required to do so (wayfinding duration; skilled navigators should be faster in reaching checkpoints and completing the levels).

**Figure 2.**
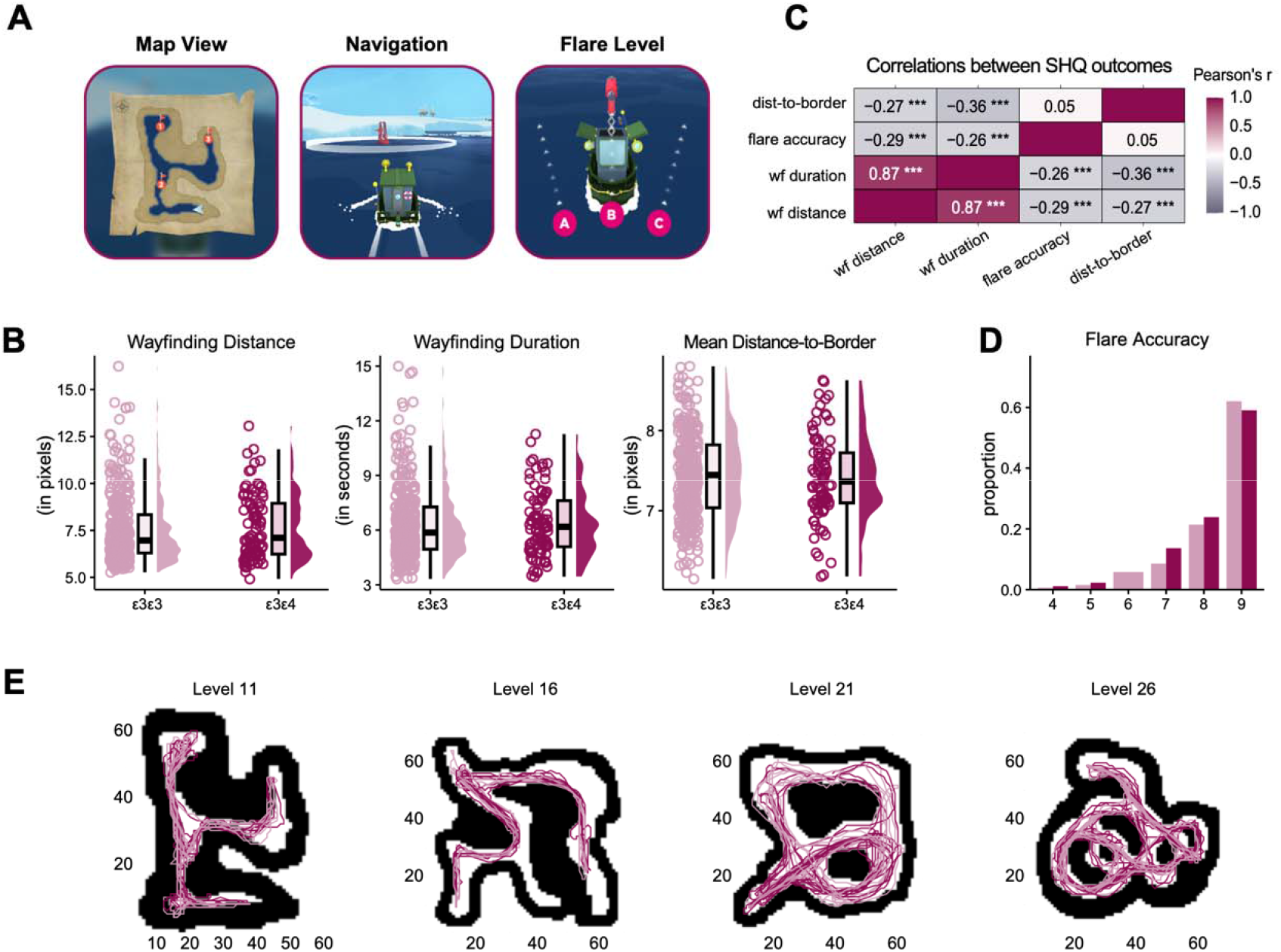
Sea Hero Quest (SHQ) game & APOE genotype effects on performance. A. Layout of the Sea Hero Quest (SHQ) mobile app showing screenshots of the allocentric view of the map with checkpoints (buoys) indicated (left panel), egocentric goal-directed spatial navigation toward a checkpoint (middle panel), and the three response options for “shooting” the flare back during a flare level (right panel). SHQ screenshots © Glitchers Limited 2026. B. Data points show individual scores on the three main wayfinding outcomes for APOE ε3ε3 carriers (in rose), and ε3ε4 carriers (in dark red). Boxplots mark the median, the interquartile range, and whiskers indicate the minimum and maximum outlier values. Group differences were assessed using Wilcoxon rank-sum tests (all p > .05). C. Heatmap illustrating correlations between the main SHQ outcomes (wf distance wayfinding distance, wf duration = wayfinding duration, dist to border = mean distance to border), with Bonferroni-corrected significance levels. D. Bar plot depicting the distribution of flare accuracy scores across all possible sum scores for the two APOE genotype groups, with bars representing the proportion of participants per score (APOE ε3ε3-carriers in rose, and APOE ε3ε4-carriers in dark red). E. Maps of the different wayfinding levels with trajectories of 10 participants per genotype group superimposed, with mean distance to border values most closely matched to their group mean.

Since gaming proficiency can shape performance on games such as the SHQ, with more experienced participants potentially performing better overall, participant scores were normalised against performance on the two practice levels (see Methods). Contrary to our expectations, results revealed no significant differences in either wayfinding distance (in pixels; n_ε3ε3_ = 327, n_ε3ε4_ = 88; mean ± SEM, ε3ε3: 7.55 ± 0.10, ε3ε4: 7.68 ± 0.19; Wilcoxon rank-sum tests, W = 13660, p = .23, r = 0.036) or wayfinding duration (in seconds; ε3ε3: 6.36 ± 0.11, ε3ε4: 6.36 ± 0.19; W = 13850, p = .29, r = 0.026) between the APOE genotype groups (Figure 2B). Results remained virtually identical after excluding extreme outliers (Supplementary Results S1). A joint measure combining wayfinding distance and duration likewise showed no significant difference between APOE genotype groups (Supplementary Results S2, Figure S1).

Lastly, goal-directed spatial navigation performance can also be characterised by how centrally participants navigated within the virtual environment, with more central routes suggesting effective route planning and confident navigation, and routes closer to the border suggesting a less efficient strategy. As ε4 carriers have previously been shown to remain closer to environmental borders while navigating [14,17], we hypothesised that young ε4 carriers in our sample would do the same. We therefore examined participants’ mean distance-to-border (see Methods), but observed no significant differences between genotype groups (in pixels; n_ε3ε3_ = 327, n_ε3ε4_ = 88; mean ± SEM, ε3ε3: 7.44 ± 0.03, ε3ε4: 7.42 ± 0.06; Wilcoxon rank-sum test, W = 14812, p = .34, r = 0.021; Figures 2C).

### No significant effects of APOE genotype on flare accuracy in the SHQ game

In addition to the wayfinding levels, the SHQ game included three “flare levels” to assess path integration, the ability to track one’s position based on self-motion cues [16]. In each flare level, participants navigated to a flare gun and then selected one of three directions (left, straight, or right) to shoot a flare back toward their starting position (Figure 2A). Flare accuracy was scored by assigning 1-3 points per level and summing across levels, with higher scores indicating better performance. While flare accuracy was previously reported to not differ between APOE ε4 carriers and non-carriers in the SHQ game [14], other studies have observed early alterations in path integration or related processes in ε4 carriers [15,17]. We therefore compared overall flare accuracy between ε3ε4 carriers and ε3ε3 carriers.

Since flare accuracy scores were discrete and skewed toward the maximum, we used Fisher’s exact test to examine the distribution of scores (range = 3-9) across genotype groups, allowing us to detect differences tied to specific score levels. Results showed no significant difference in flare accuracy between APOE genotype groups (N = 415; mean ± SEM, ε3ε3: 8.35 ± 0.06, ε3ε4: 8.34 ± 0.11; Fisher’s exact test, p = .07; Figure 2D; note that, with only three response options, 80% of participants performed at ceiling, suggesting the task may not have been sensitive enough to detect subtler group differences in path integration ability).

### SHQ performance is determined by biological sex, gaming experience, and self-reported navigation ability rather than by APOE genotype

Next, we reasoned that any effects of APOE genotype on SHQ performance might have been masked by confounding demographic (biological sex and age [36–38]), cognitive (gaming experience and self-reported navigation ability [39,40]), or task-related factors, such as map viewing duration (longer map viewing may have helped participants memorise checkpoint locations, leading to better performance). We therefore fitted linear regression models on the three main wayfinding outcomes, aggregated across levels, including APOE genotype, biological sex, age, computer- and smartphone-based gaming experience, self-reported navigation ability (SBSOD [40]), and map viewing duration as fixed effects (see Methods).

Several covariates were reliably associated with wayfinding performance (Figure 3A). Biological males outperformed females across all three outcomes, taking shorter, faster, and more central routes (wayfinding distance: p = .005, β = .15; duration: p < .001, β = .21; mean distance-to-border: p = .015, β = -.14). Higher self-reported navigation ability (SBSOD) was associated with better performance across outcomes (all p < .001; wayfinding distance: β = -.19, duration: β = -.18, mean distance-to-border: β = .17). Computer-based gaming experience predicted shorter and faster routes (wayfinding distance: p < .001, β = -.19 and duration: p < .001, β = -.21) and age was positively associated with wayfinding distance (p = .011, β = .12) and duration (p = .003, β = .14). Critically, APOE genotype remained unrelated to all three wayfinding outcomes even after covariate adjustment (all p > .05; wayfinding distance: R^2^ = .135, duration: R^2^ = .177, mean distance-to-border: R^2^ = .079). To rule out sex-specific genotype effects, we additionally tested for an APOE × biological sex interaction across the three main wayfinding outcomes, finding no evidence that genotype differentially affected navigation performance in biological males and females (all p > .05; Supplementary Results S3).

**Figure 3.**
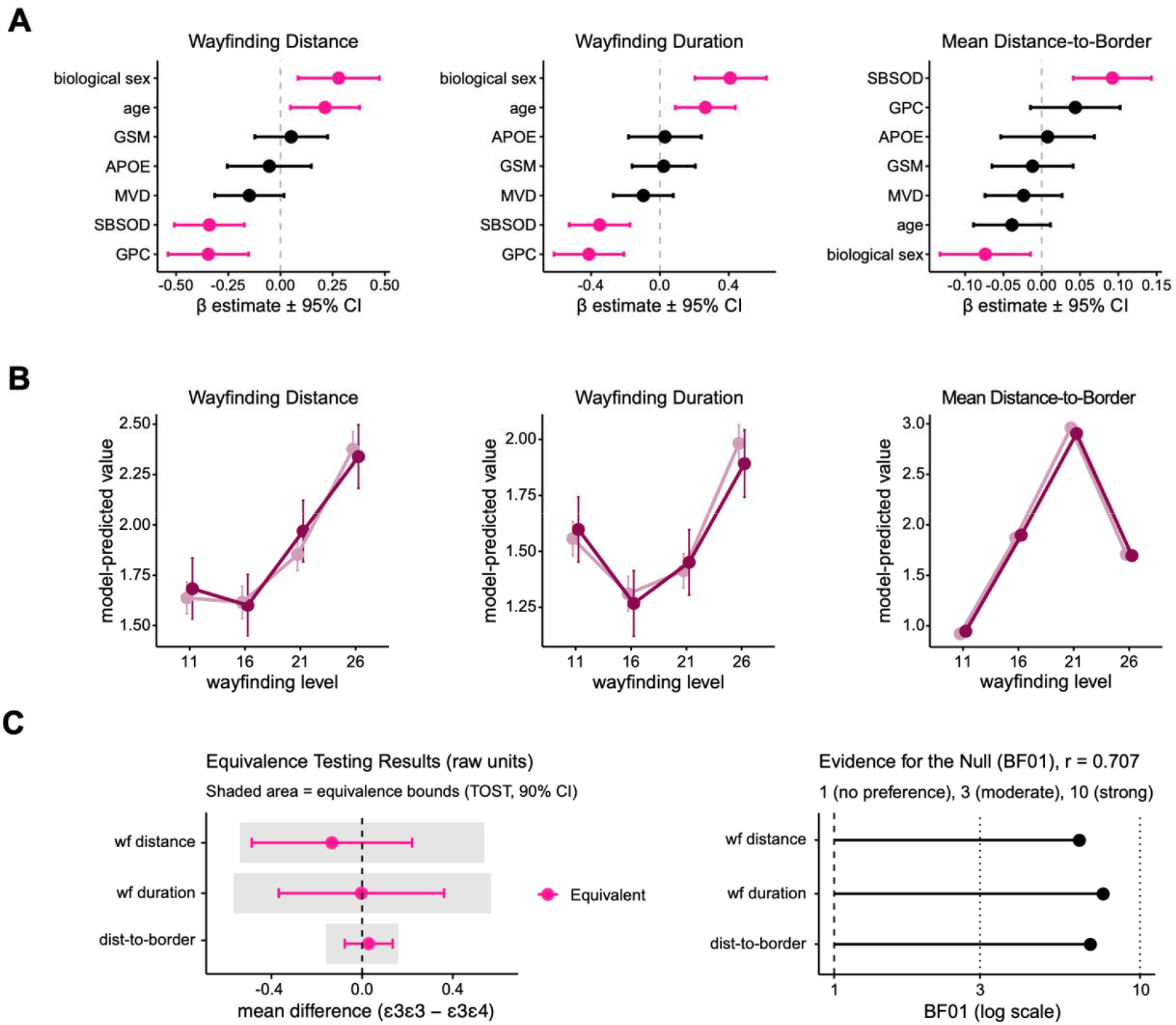
Analysis of SHQ performance measures including potentially confounding factors, learning rate, equivalence testing, and Bayes factors. A. Forest plots show the estimated regression coefficients (β-estimates ± 95% CI) for APO genotype, demographic (age, biological sex) and cognitive variables (GSM = gaming experience with navigation elements on smartphone; GPC = gaming experience with navigation elements on PC/console; MVD = map view duration; SBSOD = Sant Barbara Sense of Direction Scale) predicting wayfinding distance, wayfinding duration, and mean distance-to-border. Pink estimates and confidence intervals highlight significant predictors. B. Lines show predicted values (± 95% CI) from linear mixed-effects models for each wayfinding outcome (wayfinding distance, wayfinding duration, and mean distance-to-border) as function of APOE genotype and wayfinding level, adjusted for demographic and cognitive variables and with random intercepts for participants. APOE ε3ε3-carriers (in rose) and ε3ε4-carriers (in dark red) performed similarly across levels. C. Left panel: Mean differences between APOE genotype groups (ε3ε3/ε3ε4) for the three main wayfinding outcomes (wf = wayfinding, dist = distance) with 90% confidence intervals (corresponding to α = .05 for the two one-sided tests, TOST, equivalence test), plotted against the predefined equivalence bounds shaded in grey (± 0.3 Cohen’s d). All confidence intervals fell within these bounds, indicating statistical equivalence (marked in pink). Right panel: Bayes factors (BF) quantifying the evidence for no APO genotype effect on the three main wayfinding outcomes using a Cauchy prior on the standardised effect size (location 0, width r = 0.707). All three BF_01_ showed moderate evidence in favour of the null model.

### APOE genotype does not affect SHQ performance across levels of varying difficulty

The SHQ wayfinding levels vary in difficulty [41], and aggregating performance across levels may have concealed genotype-related differences that emerged only at particular levels [14]. To address this, we modelled level-wise performance using linear mixed-effects models, with level, APOE genotype, and their interaction as fixed effects, while controlling for demographic and cognitive covariates. Across all three wayfinding outcomes (wayfinding distance, wayfinding duration, and mean distance-to-border), performance differed significantly between levels (all χ²s(3) > 900, all p < .001). However, there was no main effect of APOE genotype (all p > .05) and no evidence for an APOE genotype × level interaction (all p > .05). Thus, accounting for systematic differences between wayfinding levels, we observed no association between APOE genotype and goal-directed spatial navigation in the SHQ game (Figure 3B).

Equivalence testing and Bayesian analyses corroborated the null effects observed for wayfinding distance, wayfinding duration, and mean distance-to-border throughout the analyses above (90% CIs within ± 0.3 Cohen’s d, all p < .05; BF_01_ = 6.34-7.58; Figure 3C; Supplementary Results S4).

### No significant effects of APOE genotype on working memory, processing speed, executive function, and face recognition

We next asked whether the APOE genotype affects cognitive abilities beyond spatial navigation, including working memory, processing speed, executive function, and face recognition, known to be impaired in AD [19–21]. Evidence for APOE genotype effects across these cognitive domains is mixed [24–27,30]. In younger adults, some studies proposed a transient cognitive advantage in young APOE ε4 carriers, with detrimental effects emerging only later in life, consistent with an antagonistic pleiotropy account [27,31,42]. However, reported effects appear to depend on additional factors, such as education level [43] or task difficulty, with ε4 carriers performing better on tasks requiring higher cognitive control [31]. In older adults, ε4 carriers performed less well on tasks requiring learning and verbal recall [44], attentional shifting [7], and working memory [29]. Overall, APOE ε4 effects appear subtle and highly selective across cognitive domains, particularly in young adults [27,28]. We therefore tested for potential APOE genotype effects on working memory, executive function, and face recognition in healthy young adults using four tasks designed to probe these cognitive domains (Figure 4A).

**Figure 4.**
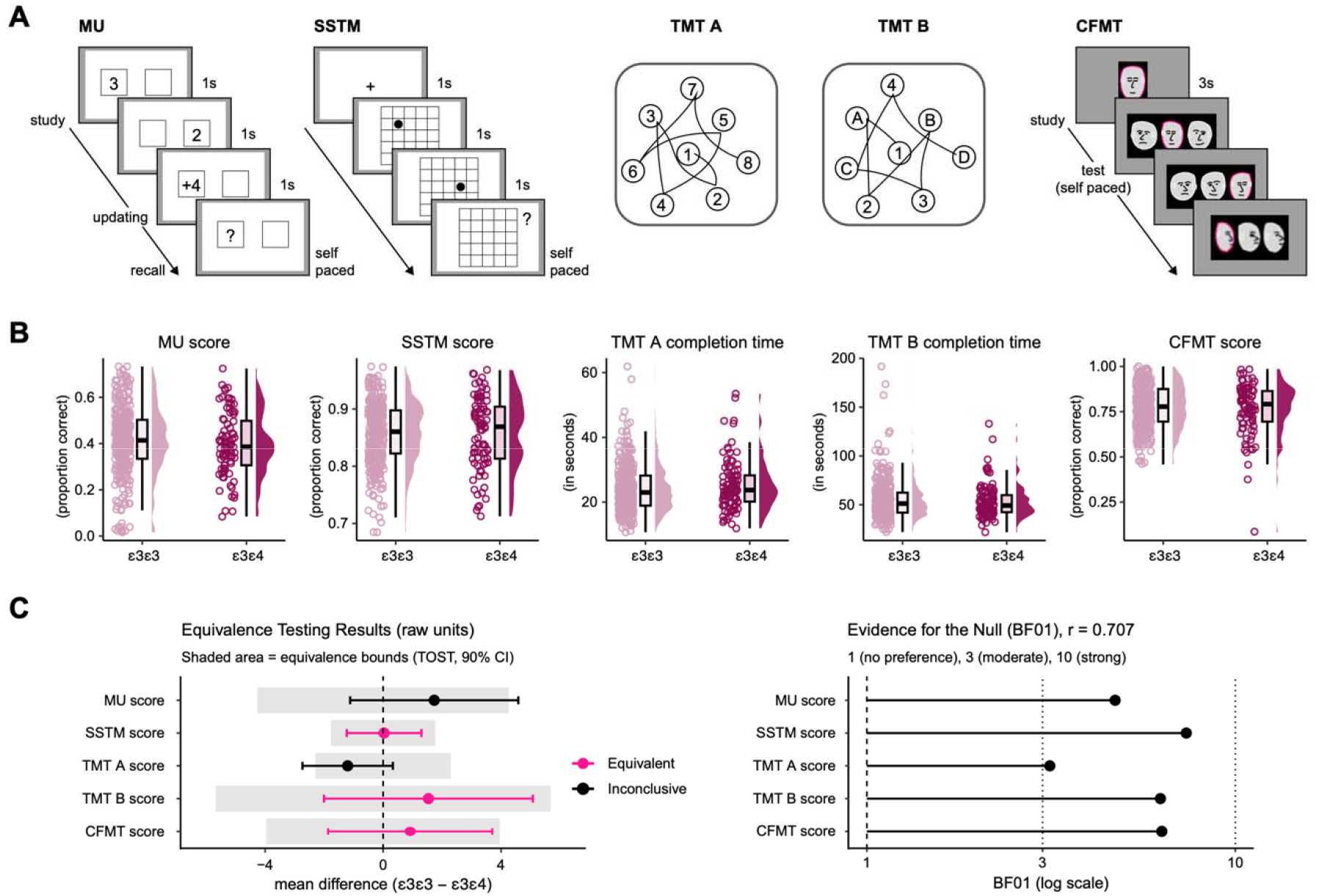
Tasks assessing working memory, executive function, and face recognition & APOE genotype effects on performance, including equivalence testing and Bayes factors. A. Task layout for the five additional cognitive tasks: Th Memory Updating (MU) task prompted participants to memorise and update numbers. In the Spatial Short-Term Memory (SSTM) task, they viewed and reproduced dot patterns. In the Trail Making Test, participants connected numbers (TMT A) and both numbers and letters (TMT B) in ascending order. The Cambridge Face Memory Test (CFMT) assessed their ability to recognise faces using photographs. In the schematic illustration the target face is marked in pink. B. Data points show individual scores on all cognitive tasks for APOE ε3ε3 carriers (in rose), and APOE ε3ε4 carriers (in dark red). Boxplots mark the median, the interquartile range, and whiskers indicate the minimum and maximum outlier values. Group differences were assessed using Wilcoxon rank-sum tests (all p > .05). C. Left panel: Mean differences between APOE genotype groups (ε3ε3/ε3ε4) for all cognitive task measures with 90% confidence intervals (corresponding to α = .05 for the two one-sided tests, TOST, equivalence test), plotted against the predefined equivalence bounds shaded in grey (± 0.3 Cohen’s d). Confidence intervals of SSTM, CFMT, and TMT B lay within equivalence bounds, indicating statistical equivalence (marked in pink). MU and TMT confidence intervals exceeded the bounds (marked in black). Right panel: Bayes factors (BF) quantifying the evidence for no APOE genotype effect on all cognitive task measures using a Cauchy prior on the standardised effect size (location 0, width r = 0.707). All BF_01_ showed moderate evidence in favour of the null model.

To assess working memory, we used a memory updating (MU) task in which participants memorised digits and continuously updated them following a series of arithmetic operations, and a spatial short-term memory (SSTM) task in which participants reproduced sequences of dots presented on a grid [45] (see Figure 4A and Methods). MU performance, quantified as the proportion of correctly recalled digits, did not differ significantly between ε3ε3 and ε3ε4 carriers (N = 392; mean ± SEM, ε3ε3: 0.414 ± 0.0083, ε3ε4: 0.397 ± 0.0151; Wilcoxon rank-sum test, W = 14186, p = .91, r = 0.069). Similarly, SSTM performance, quantified as the proportion of correctly reproduced dot patterns, did not differ between genotype groups (N = 389; mean ± SEM, ε3ε3: 0.858 ± 0.0031, ε3ε4: 0.857 ± 0.0070; Wilcoxon rank-sum test, W = 12532, p = .43, r = 0.009).

To test whether visual search, processing speed, mental flexibility, and executive functions varied between APOE genotype groups, we next evaluated the Trail Making Test (TMT A & B) [46,47] in which participants connected numbers (TMT A) or numbers and letters in ascending numerical and alphabetical order (TMT B; see Figure 4A and Methods). Performance was measured by completion time, with shorter completion time indicating better performance. We observed no significant APOE genotype effects on TMT A (N = 427; mean ± SEM, ε3ε3: 24.1 ± 0.40, ε3ε4: 25.3 ± 0.84; Wilcoxon rank-sum test, W = 14112, p = .11, r = 0.060) or TMT B performance (N = 427; mean ± SEM, ε3ε3: 55.1 ± 1.10, ε3ε4: 53.5 ± 1.84; Wilcoxon rank-sum test, W = 15997, p = .71, r = 0.027).

Last, we examined whether potential cognitive differences extended to face recognition. Participants completed the Cambridge Face Memory Test (CFMT) [48] that assesses recognition of previously unfamiliar faces among novel faces (see Figure 4A and Methods). CFMT performance did not differ significantly between APOE genotype groups (N = 406; mean ± SEM, ε3ε3: 0.774 ± 0.0066, ε3ε4: 0.765 ± 0.0154; Wilcoxon rank-sum test, W = 13875, p = .45, r = 0.006) and was in the normal, previously reported range for healthy young adults (aged 18-26, mean ± SEM = 0.804 ± 0.0156) [48]. Together, these findings suggest that ε3ε3 and ε3ε4 carriers performed similarly across tasks probing working memory, executive function, and face recognition (Figure 4B).

Equivalence testing and Bayesian analyses corroborated the null effects for SSTM, TMT B, and CFMT (90% CIs within ± 0.3 Cohen’s d, all p < .05; BF_01_ = 6.32-7.37; Figure 4C). For MU and TMT A, equivalence bounds were slightly exceeded, and Bayes factors showed weaker support for the null (BF_01_ = 3.14-4.72), suggesting that any true performance differences are likely minimal (Supplementary Results S5).

### Exploratory analysis of other APOE genotypes: homozygous APOE ε4 carriers and APOE ε2 carriers

AD risk increases with the number of APOE ε4 alleles, from an approximately 2-3-fold increase with one ε4 allele to a 10-15-fold increase with two alleles [6,33,35]. Also, cognitive differences associated with APOE ε4 appear more pronounced in homozygous carriers [27,49,50]. We had not observed any significant differences between ε3ε3 and ε3ε4 carriers, potentially suggesting that the effects of a single ε4 allele may be too subtle to detect. We thus reasoned that performance differences may become apparent when comparing ε3ε3 and ε4ε4 carriers. ε4 homozygotes are particularly rare [35] and thus accounted for a small fraction of our sample (11 ε4ε4 carriers among the 1,000 genotyped individuals). Similarly rare and less studied is the APOE ε2 allele. It was previously linked to reduced AD risk and potential protective effects on cognition [51], but whether ε2-related differences in behavioural performance can already be detected in healthy young adults remains unclear. To explore potential differences between ε3ε3 carriers, ε4 homozygotes and ε2 carriers (ε2ε2, ε2ε3, and ε2ε4), we used a non-parametric bootstrapping procedure and repeatedly resampled participants within each genotype group (to account for the small sample sizes; 10,000 resamples per contrast; see Methods).

We first focused on goal-directed spatial navigation and examined the three main wayfinding outcomes (wayfinding distance, wayfinding duration, and mean distance-to-border), comparing ε3ε3 carriers against each other genotype group (ε3ε4, ε4ε4, ε2ε2, ε2ε3, ε2ε4). For wayfinding distance and duration, all confidence intervals included zero, indicating no genotype-related differences. For mean distance-to-border, however, ε2ε3 carriers stayed closer to the borders than ε3ε3 carriers, possibly reflecting greater reliance on environmental boundaries and reduced confidence in their navigational strategy (Δ = 0.16, 95% CI [0.01, 0.31], not corrected for multiple comparisons; Figure 5A).

**Figure 5.**
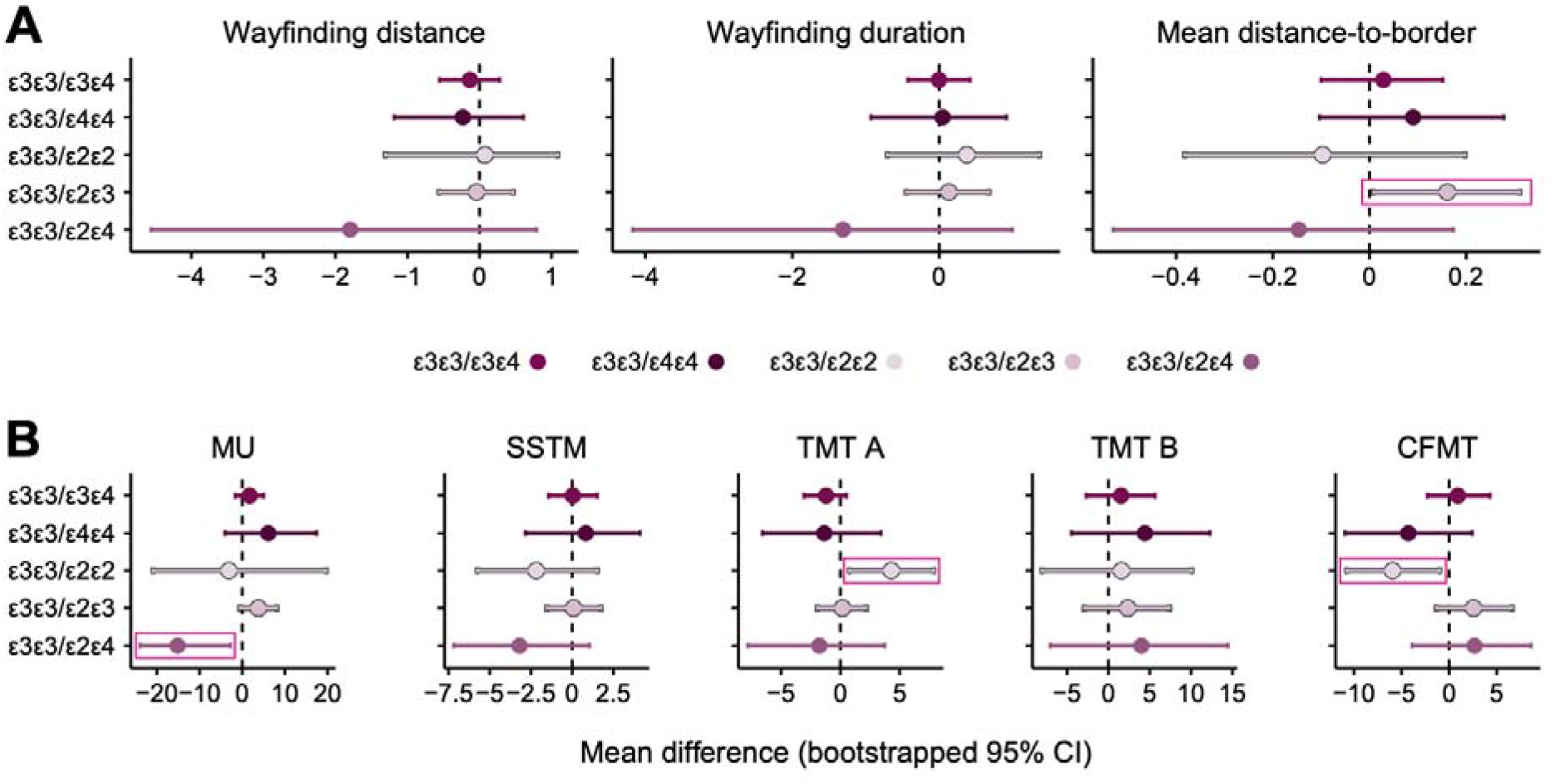
Bootstrapped APOE genotype comparisons for spatial navigation and cognitive tasks assessing working memory, executive function, and face recognition. A. Bootstrapped mean differences (10,000 resamples per contrast) between APO genotype groups for the three main wayfinding outcomes: wayfinding distance, wayfinding duration, and mean distance-to-border. B. Bootstrapped mean differences between APOE genotype groups for MU, SSTM, TMT A & B, and CFMT performance. Points represent estimated mean differences between genotype groups (ε3ε3/ε3ε4, ε3ε3/ε4ε4, ε3ε3/ε2ε2, ε3ε3/ε2ε3, ε3ε3/, ε2ε4 ), with vertical lines indicating 95% confidence intervals. Comparisons for which the confidence intervals do not include zero are marked in pink, indicating a performance difference between groups. Performance differences were not observed between ε3ε3 carriers and ε4 homozygotes, but were found in ε2 carriers, who navigated closer to the environmental boundaries (mean distance-to-border) and performed better on MU, TMT A, and CFMT.

We then examined performance across the remaining cognitive tasks (MU, SSTM, TMT A & B, CFMT). For MU, SSTM, and TMT B, all genotype contrasts yielded confidence intervals that included zero, indicating no significant genotype-related differences. However, ε2ε4 carriers showed better memory updating on the MU (Δ = -15.14, 95% CI [-23.89, -2.30]), ε2ε2 carriers were faster to complete the TMT A, and recognised more faces on the CFMT than ε3ε3 carriers (TMT A: Δ = 4.29, 95% CI [0.72, 7.93]; CFMT: Δ = -5.95, 95% CI [- 10.83, -1.02]; all not corrected for multiple comparisons; Figure 5B).

Overall, we observed no performance differences between ε3ε3 and ε4 homozygote carriers, while ε2 carriers navigated closer to environmental borders and outperformed ε3ε3 carriers on memory updating, processing speed, and face recognition, consistent with reports of protective effects.

## Discussion

In the present study, we investigated whether carrying the APOE ε4 allele, which represents the major genetic risk factor for AD, is associated with differences in spatial navigation and broader cognition in healthy young adults. Across a large sample, APOE ε3ε3 and ε3ε4 carriers performed comparably on spatial navigation (SHQ), working memory, executive functioning, and face recognition, with converging evidence from equivalence testing and Bayesian analyses corroborating these null effects. Exploratory analyses based on small subgroups across all genotypes, however, suggested a more differentiated pattern involving the less-studied APOE ε2 allele. ε2ε3 carriers navigated closer to environmental borders in the SHQ game, ε2ε4 carriers showed better memory updating, and ε2ε2 carriers showed faster processing speed and better face recognition. Together, these findings offer new insights into the underexplored role of APOE ε2 in young adults, suggesting that while APOE ε4 showed no detectable effects among healthy young adults, the ε2 allele may confer cognitive benefits.

We hypothesised that spatial navigation performance in the SHQ game would be compromised in young adult ε4 carriers (18-35 years), who are at increased genetic risk for AD. Contrary to this prediction, we observed no significant performance differences between ε3ε3 and ε3ε4 carriers. Both completed the wayfinding levels with comparable efficiency, navigated at similar distances from the borders, and showed no significant differences in path integration, a pattern that held across SHQ levels of varying difficulty. This may appear to contradict previous work showing that SHQ performance can distinguish ε4 carriers from non-carriers in middle-aged and older adults [14], yet aligns with evidence from younger cohorts. For instance, Bierbrauer et al. [15] found no consistent association between ε4 status and path integration in younger adults (18-41 years), and Kunz et al. [17] reported intact navigation performance in young ε4 carriers (18-31 years), despite alterations at the neural level. Together, these results indicate that age is a key factor in ε4-related effects, with spatial navigation remaining largely intact in younger ε4 carriers and deficits potentially emerging only later in life.

We next asked whether ε4-related effects extended to other cognitive domains commonly impaired in AD [19–21], and found no genotype-related differences in working memory, executive functioning, and face recognition. This is broadly consistent with prior work in younger adults, reporting only negligible ε4 effects in mid-adulthood (35-60 years) [30] and no significant effects in younger ε4 carriers across cognitive domains (9-31 years) [25]. As reviewed by Daly et al. [27], evidence for ε4-related cognitive effects remains mixed and often depends on the cognitive domain and task at hand. Overall, we found that any ε4-related effects on cognitive performance appear minimal in young adulthood.

As a final exploratory step, we focused on ε4 homozygotes, reasoning that ε4-related cognitive differences may be most pronounced in carriers of two ε4 alleles, who are at markedly higher AD risk [27,49,50]. Yet ε4 homozygotes did not significantly differ from other genotype groups on any of the behavioural measures assessed. Although these findings should be interpreted cautiously, given the small number of homozygous individuals in our sample, they further support the notion that ε4-related behavioural effects in young adults are likely very subtle and difficult to detect with these common cognitive-behavioural approaches.

Even though our results suggest preserved behavioural performance in healthy young ε4 carriers, early genotype-related changes may be masked by compensatory neural mechanisms before becoming behaviourally observable in older age. Bookheimer et al. [52] reported higher brain activation during memory encoding and recall in ε3ε4 carriers relative to ε3ε3 carriers, suggesting that ε4 carriers recruit additional brain regions to maintain performance. Functional neuroimaging in our sample may therefore reveal activation differences that are not yet visible at the behavioural level. ε4-related effects likely also depend on additional risk factors such as vascular burden, inflammation, or family history of dementia [53,54]. Newton et al. [54], for instance, found poorer path integration specifically among ε4 carriers with a parental history of dementia, indicating that such effects may only emerge alongside additional vulnerabilities absent from our otherwise healthy young sample. Future work should examine how genetic and other risk factors jointly shape cognitive performance.

Another possibility is that our tasks lacked the sensitivity to detect early ε4-related change in such a young sample. Spatial navigation is considered a promising marker of early AD-related impairment, but its sensitivity varies considerably depending on the particular paradigm used [27]. The SHQ wayfinding levels in this study were relatively easy [41], which may explain the absence of genotype effects. Path integration, the ability to update one’s position based on self-movement cues, may offer a more sensitive measure. It relies on entorhinal circuits affected early in AD [12] and is impaired in individuals at increased AD risk [13,15]. The SHQ flare levels were designed to probe path integration directly, requiring participants to select one of three directions to shoot a flare back to their starting point after navigating away from it. We observed no genotype differences here, and performance was near ceiling. However, the environment remained visible throughout, so landmark and boundary cues were still available. The task, therefore, did not isolate path integration based on self-movement alone and may have missed subtle impairments [15]. Moreover, choosing between three discrete response options may have been too easy for this young, high-functioning sample. This pattern is consistent with Coughlan et al. [14], who reported no differences on the same flare levels even in an older, higher-risk sample. We encourage future research to use path integration measures that capture performance more precisely and rely less on landmark and boundary information, to clarify whether subtle ε4-related differences are already present in young adults.

Alternatively, spatial navigation paradigms that involve active movement through physical or virtual space may offer additional insights. Such navigation tasks combine the cognitive aspects of navigation with the sensorimotor processes supporting goal-directed movement, which may already be affected in at-risk individuals [55,56]. They may capture aspects of navigation less evident in stationary, screen-based formats and could be useful for studying subtle differences in navigation. However, virtual, screen-based navigation remains a valid approach. SHQ performance has been shown to predict real-world spatial navigation performance [57] and self-reported navigation ability (SBSOD) predicted SHQ wayfinding performance in our sample (all p < .001), with better self-rated navigators taking shorter, faster and more central routes.

While we found no evidence linking ε4 to behavioural performance, exploratory analyses across all genotype groups indicated ε2-related differences in cognition. ε2ε3 carriers navigated closer to the virtual borders in the SHQ game, ε2ε4 carriers performed better on memory updating, and ε2 homozygotes showed faster processing speed and better face recognition. The ε2 allele is less common than ε4 and has therefore received comparatively little attention, despite evidence of protective effects in older adults, including lower AD risk, later disease onset, longevity, reduced neuropathology, and slower cognitive decline [32–34,58]. It has also been linked to differences in medial temporal lobe structure, including greater hippocampal grey matter volume in young and older adults [59,60] and greater entorhinal cortical thickness in childhood [61]. During virtual navigation, ε2 carriers also showed stronger hippocampal involvement than ε3 and ε4 carriers, suggesting that ε2 may shape how spatial information is processed [60]. Our findings extend this work and point to a cognitive advantage even at a young age, though the effects varied across genotype groups and tasks and were based on a very small ε2 subsample, warranting replication in larger samples.

To conclude, we examined how ε4 relates to spatial navigation and broader cognition in healthy young adults. By combining several cognitive tasks with complementary analytical approaches, we provide a careful test of whether ε4-related differences are already visible earlier in life. We found no evidence that ε3ε4 carriers performed worse than ε3ε3 controls. Exploratory analyses of other genotypes instead pointed to distinct patterns in ε2 carriers, consistent with prior work linking ε2 to reduced AD risk and possible protective effects. Overall, our findings suggest that APOE-related cognitive differences in young adults are likely subtle and task-dependent, highlighting the challenge of detecting early cognitive signatures of genetic dementia risk.

## Methods

### Sample

A total of 1,000 individuals enrolled in the study. For the SHQ, 108 individuals were excluded for not completing the study protocol (questionnaires and/or SHQ), 393 were excluded due to self-reported neurological or psychiatric conditions (such as epilepsy, migraine, major depressive, anxiety, or eating disorders), 76 were excluded because they did not carry the required genotype (APOE ε3ε3 or ε3ε4), and 8 were excluded for not completing all SHQ task levels. The final SHQ sample thus comprised 415 participants (APOE ε3ε3: 200 female, 127 male, age range 18-34 years; APOE ε3ε4: 62 female, 26 male, age range 18-35 years). Additional demographic information is provided in Table 1, and the exclusion procedure is summarised in Figure 1.

For the remaining cognitive tasks, sample sizes varied slightly because not all participants completed all study components. This was partly due to a temporary license suspension that prevented access to the three tasks administered via the online platform Labvanced (https://www.labvanced.com). In total, 94 participants were excluded for not completing the protocol (questionnaires), 399 for self-reported neurological or psychiatric conditions, and 80 for not carrying the required genotype. Participants without valid task responses (scores of 0) were additionally excluded, leaving the following samples for analysis (APOE ε3ε3/ε3ε4): Memory Updating Task: n = 308/84, Spatial Short-Term Memory Task: n = 311/83, Cambridge Face Memory Test: n = 318/88, Trail Making Test A & B: n = 335/92. The exclusion procedure is summarised in Figure 1.

The SHQ and cognitive task samples were largely overlapping. All SHQ participants were included in the TMT A & B analyses, with an additional 12 participants contributing TMT data only. Overlap for the remaining tasks was slightly reduced due to missing data. All results were robust to alternative preprocessing choices, including using the full sample and excluding outliers beyond ±3 median absolute deviations (MAD) within each score variable. All participants included in the final sample were healthy, reported no history of neurological and/or psychiatric disorders, and provided written informed consent prior to participation. Participants received either monetary compensation or course credit. The study protocol was reviewed and approved by the local ethics committee of the University of Vienna (reference number 00538).

### Procedure

This study served as a prescreening to create a participant pool with two groups based on genetic risk for Alzheimer’s Disease (AD), one group with a genotype linked to increased risk (APOE ε3ε4-carriers) and one with no such risk (APOE ε3ε3-carriers), with the aim of reinviting eligible participants to take part in follow-up studies (not further discussed here). The study involved two sessions held on consecutive days. On Day 1 (onsite in the lab), participants had a DNA sample taken and performed the Trail Making Test (TMT A & B). On Day 2 (online), participants completed the Sea Hero Quest (SHQ), an app-based game to assess spatial navigation ability, three computer-based tasks testing working memory (Memory Updating, MU; Spatial Short Term Memory, SSTM) and general face recognition ability (Cambridge Face Memory Test, CFMT), as well as a set of questionnaires. All tasks and questionnaires that were included in our analyses are described in detail below.

#### Trail Making Test (TMT A & B)

The Trail Making Test (TMT) [46,47] is a well-established neuropsychological paper-and-pencil test that is frequently included in neuropsychological batteries assessing AD [62]. The TMT consists of two parts and measures visual search, scanning, processing speed, mental flexibility, and executive functions. In TMT A, participants were required to connect 25 numbers in ascending order. In TMT B, participants alternately connected 25 numbers and letters in ascending numeric and alphabetical order. Participants were instructed to complete both parts as quickly as possible, without lifting the pen from the paper, while completing the task. Performance on both parts was quantified as the completion time in seconds.

#### Sea Hero Quest (SHQ)

Spatial navigation ability was assessed using the Sea Hero Quest (SHQ; www.seaheroquest.com [63]), an app-based game developed in 2015 by researchers at University College London (UCL) and the University of East Anglia in collaboration with the game developer GLITCHERS, with support from Deutsche Telekom, as a tool to measure spatial navigation performance. The game is available for Android and iOS devices, and access was provided via individual participant keys. Participants completed the task remotely on their own devices following standardised instructions.

After completing two mandatory practice levels (levels 1 and 2), participants proceeded to two types of navigation levels: goal-directed “wayfinding levels” (levels 11, 16, 21, and 26) and “flare accuracy levels” (levels 4, 14, and 24). In wayfinding levels, participants first viewed an allocentric map indicating the starting location and a set of checkpoints to be visited in a predefined order. Once the map disappeared, participants navigated a boat through virtual river landscapes that varied across levels to reach the checkpoints as quickly and accurately as possible. The boat was controlled by right/left taps for steering and upward swipes to temporarily increase speed. Navigation relied on self-motion cues and environmental features. In flare accuracy levels, no allocentric map was provided. Participants navigated the boat through the virtual environment to locate a flare gun. Upon reaching the flare gun, the boat was rotated by 180°, and participants were required to fire the flare back toward their starting location by selecting one of three response options (left/A, front/B, or right/C). For both level types, participants received one to three stars as feedback on their performance.

Spatial navigation performance during wayfinding levels was quantified using two primary outcome measures: wayfinding distance and wayfinding duration. Wayfinding distance was defined as the cumulative path length travelled by participants during a level, computed by summing Euclidean distances between successive recorded x/y coordinates. Wayfinding duration was quantified as the total time (in seconds) required to complete a level. To account for individual differences in general gaming proficiency, wayfinding distance and duration in each test level were normalised within participants using performance in the two training levels (Levels 1-2). Specifically, wayfinding distance and duration in each test level were divided by participants’ summed distance and summed duration across the two training levels, respectively. Wayfinding distance and duration are strongly related and capture overlapping aspects of navigation efficiency (shorter paths typically result in shorter completion times). However, in some cases, path length and navigation speed may not align. Some participants may have needed more decision time and therefore taken longer to complete a short path, or moved quickly on a longer, less direct route. Thus, we additionally derived a composite wayfinding performance score by summing the z-standardised distance and duration measures and multiplying the result by -0.5, such that higher values reflected shorter path lengths and completion times. We further quantified participants’ mean distance-to-border during wayfinding levels based on the recorded x/y coordinates of their navigation trajectories. For each level, we specified a map indicating the distance of each location within the virtual environment to the nearest environmental boundary. Using participants’ x/y coordinates, we determined their distance to the border at each time point by referencing this map. These distances were then averaged across the trajectory to obtain a mean distance-to-border value for each level, capturing whether participants tended to navigate close to the borders or further toward the centre of the virtual environment, with higher values indicating trajectories that deviated further from the border, including more central routes.

Flare accuracy served as an index of path integration, reflecting how well participants kept track of their starting position while moving through the environment. Performance in flare accuracy levels was quantified based on the direction (left /A, front/B, or right/C) participants indicated back to their starting position following the 180° rotation. Responses were scored on a three-level scale corresponding to one (low accuracy), two (medium accuracy), or three points (high accuracy) per level. Flare accuracy scores were summed up across the three flare accuracy levels to obtain individual total scores, with possible values ranging from 3 to 9.

#### Memory Updating (MU)

Memory updating ability was assessed using an adapted version of the Memory Updating task (MU) [45], implemented in Labvanced (https://www.labvanced.com) to allow for online administration. In each trial, participants were presented with a set of 3-5 frames containing digits to be memorised. Initial digits appeared sequentially for 1 s each. Following encoding, 2-6 arithmetic operations (e.g., “+2” or “-4”) were displayed sequentially for 1 s each, pertaining to one frame at a time. Participants were instructed to apply each operation to the digit currently held in memory and to update the memorised value accordingly. Final recall was prompted by question marks appearing sequentially in each frame. Responses were not time-limited. The task consisted of two practice trials and 15 test trials. Arithmetic operations ranged from +7 to -7. Not every frame was necessarily updated within a trial, and the same frame could be updated multiple times. Task performance was quantified as the average proportion of digits that were correctly updated and recalled across trials, yielding a single performance index per participant.

#### Spatial Short Term Memory (SSTM)

Spatial working memory was assessed using an adapted version of the Spatial Short-Term Memory task (SSTM) [45], implemented in Labvanced to allow for online administration. In each trial, participants first viewed a fixation cross (1 s), followed by a sequence of 2-6 dots presented one at a time (1 s) at random locations on a 10 × 10 grid displayed on the computer screen. After stimulus presentation, participants were asked to reproduce the dot pattern in a self-chosen order by clicking on the corresponding locations on an empty grid. Responses could be corrected prior to final submission, and there was no time limit for doing so. The task included two practice trials and 30 test trials. On each trial, dots were presented either across the entire 10 × 10 grid or restricted to a single quadrant (5 × 5), with both display conditions occurring with equal probability. Task performance was quantified as the similarity between the presented and reproduced dot patterns, independent of response order and absolute location. For scoring, the presented dot pattern was systematically shifted as a whole to best align with the reproduced pattern, such that overall displacement of the pattern did not affect the score. Participants received 2 points for each dot recalled at the exact location, 1 point if the recalled dot deviated by one grid position from the target location, and 0 points if the deviation exceeded one grid position. Trial-wise scores were converted into proportional scores ranging from 0 to 1. Proportional scores were then averaged across trials to obtain a single performance index per participant.

#### Cambridge Face Memory Test (CFMT)

We assessed general face recognition ability using the Cambridge Face Memory Test (CFMT) [48], administered via Labvanced. Participants studied 6 different male faces from left-, frontal-, and right- viewed photographs. All photographs were cropped to display only the facial region. Each face stimulus was presented for 3 s, followed by an inter-trial interval (ITI) of 500 ms. Recognition memory was tested in three stages. Before the onset of the second and third stages, participants reviewed all six previously learned faces presented together for 20 s. First, participants were asked to distinguish the previously studied target faces from two distractors (drawn from a fixed pool of 46 distractor faces that were repeated across trials). Second, the target faces were displayed from novel viewpoints, and third, target faces were additionally degraded with visual noise. Performance was quantified as the proportion of correctly recognised faces across all trials.

#### Santa Barbara Sense of Direction Scale (SBSOD)

Self-reported spatial navigation ability was assessed using the Santa Barbara Sense of Direction Scale (SBSOD) [40]. The SBSOD is a widely used questionnaire that measures individuals’ perceived sense of direction and everyday navigational competence. It consists of a single scale comprising 15 items, which assess abilities such as wayfinding, use of spatial cues, and orientation in familiar and unfamiliar environments. Participants rated each item on a 7-point Likert scale (1 = “strongly agree; 7 = “strongly disagree”), with higher scores indicating a better self-reported sense of direction. Positively worded items were reverse-scored prior to analysis, and item scores were averaged to yield a single SBSOD score per participant. The SBSOD has good internal consistency (Cronbach’s α = 0.88) and high test-retest reliability (r = 0.90).

### DNA sample collection and APOE genotyping

Buccal swab samples were collected with sterile cotton swabs (Sarstedt AG & Co. KG, Nümbrecht, Germany) from both inner cheeks. Samples were stored at room temperature or, if kept for longer than a week, at -25 °C until DNA extraction using the SwabSolution™ Kit (Promega, Madison, USA) according to the manufacturer’s instructions.

APOE genotyping was performed applying TaqMan-probe-based quantitative real-time PCR (qPCR) based on a modified version of a published protocol [64]. Briefly, APOE genotypes were determined by differential amplification with three specific reaction setups for the ε2-, ε3- and ε4 alleles, respectively. 20 μL reaction mixes were prepared containing 1× Luna® Universal Probe qPCR Master Mix (New England Biolabs, Ipswich, MA, USA), 2 μL 5× AmpSolution™ Reagent (Promega), 0.5 μM of an allele-specific APOE primer/probe combination, 0.5 μM ACTB (beta-actin) primer and probe, and 2 μL DNA extract. Detailed sequence information of primers and probes is provided in Supplementary Table 1. No-template controls were included for each reaction mix in each run. For each sample, all three analyses were performed simultaneously on the same 96-well PCR plate (Thermo Fisher Scientific, Waltham, MA, USA) to reduce possible inter-run variations. The following thermal protocol was applied using the Applied Biosystems™ QuantStudio™ 5 Real-Time PCR System (Thermo Fisher Scientific): enzyme activation (hot start) at 95 °C for 1 min, followed by 40 cycles with denaturation at 95 °C for 15 s, and annealing/extension at 65 °C for 30 s. The APOE genotype was determined by the ΔCt approach, i.e. calculated by subtracting the Ct value of ACTB from the Ct values of the three different APOE alleles (ε2, ε3 and ε4). The ΔCt values ≤ 5 were considered as positive and ΔCt values ≥ 10 were considered as negative for the ε2-, ε3- and ε4 alleles, respectively.

Genotyping results were confirmed for selected samples by direct sequencing, i.e., PCR amplification of the APOE gene region containing the allele-discriminating single nucleotide polymorphisms (SNPs) rs429358 and rs7412, followed by Sanger sequencing using the BigDye™ Terminator v3.1 Cycle Sequencing Kit (Thermo Fisher Scientific) and electrophoretic separation on the AB 3500 Genetic Analyzer (Thermo Fisher Scientific).

APOE genotyping yielded the following allele combinations: ε2ε2, ε2ε3, ε2ε4, ε3ε3, ε3ε4, and ε4ε4. Since the ε3ε3 and ε3ε4 genotypes are the most prevalent in the population [65], the main analyses for this study focused on comparisons between these two genotypes.

Data collection was double-blinded. As there are no clear treatment recommendations for the assessed APOE genotypes, and their predictive value for individual development is limited, APOE genotype information was not disclosed to participants (similar to previous studies in the field [15,17]).

### Data preprocessing and statistical analysis

Data were first preprocessed in R (version 4.2) for the SHQ and in MATLAB (2018a) for all other online tasks, and then analysed in R. Duplicate subject entries were removed during preprocessing to ensure one valid record per participant. Demographic characteristics of APOE ε3ε3 carriers and ε3ε4 carriers were compared using two-tailed independent samples t-tests (t.test in R) for age and χ^2^ tests (chisq.test in R) for biological sex and education.

We next examined group differences in task performance between APOE ε3ε3 carriers and ε3ε4 carriers using one-tailed Wilcoxon rank-sum tests to account for non-normal data distributions. For the SHQ, performance was normalised prior to statistical analysis by dividing level-wise performance by the summed score of the two practice levels, as described previously [14], to control for individual differences in player proficiency. Bonferroni correction was applied to account for multiple comparisons. Unless stated otherwise, effect sizes are reported as Cohen’s d.

To examine the effects of APOE genotype on performance in SHQ wayfinding levels while accounting for demographic, cognitive, and task-related factors, we fitted linear regression models in R using participants’ mean performance across levels. Models were fitted separately for the three main wayfinding outcomes: wayfinding distance, wayfinding duration, and mean distance-to-border. We included APOE genotype, biological sex, age, computer- and smartphone-based gaming experience involving virtual navigation, self-reported navigation ability (as measured by the SBSOD), and map viewing duration as fixed effects.

To test for a potential APOE genotype × biological sex interaction, we specified additional linear regression models fitted separately for each wayfinding outcome (wayfinding distance, wayfinding duration, and mean distance-to-border). To assess whether demographic, cognitive, and task-related covariates altered a potential APOE genotype × biological sex interaction, we fitted covariate-adjusted linear regression models that included the APOE genotype × biological sex interaction, as well as age, computer- and smartphone-based gaming experience involving virtual navigation, self-reported navigation ability (SBSOD), and map viewing duration as fixed effects.

To examine whether APOE genotype effects on navigation performance varied across wayfinding levels of differing game characteristics, we analysed level-wise performance data using linear mixed-effects models implemented in the nlme package in R. These models were fitted separately for each wayfinding outcome (wayfinding distance, wayfinding duration, and mean distance-to-border) and included level (11, 16, 21, 26), APOE genotype, and their interaction as fixed effects, alongside demographic, cognitive, and task-related covariates (age, computer- and smartphone-based gaming experience involving virtual navigation, self-reported navigation ability (SBSOD), and map viewing duration). Random intercepts for participants were included to account for the dependence of repeated observations within individuals.

Linear regression models were fitted using ordinary least squares estimation, and linear mixed-effects models were fitted using restricted maximum likelihood (REML). Model assumptions were assessed using graphical diagnostics, including the inspection of standardised residuals plotted against fitted values and residual Q-Q plots (Supplementary Methods S2). Unless stated otherwise, all statistical tests were two-tailed with α = .05.

Deviations from the preregistered analysis plan are reported in Supplementary Methods S1.

### Equivalence testing and Bayes factors

To determine whether the absence of APOE genotype effects reflected statistical equivalence rather than insufficient power, we applied equivalence testing and Bayesian hypothesis testing to the main wayfinding outcomes of the SHQ and the other cognitive task measures.

Equivalence testing was performed using the two one-sided tests (TOST) procedure [66,67]. Standardised equivalence bounds were defined a priori at ± 0.3 Cohen’s d. For each outcome measure, the standardised mean difference between APOE ε3ε3 and ε3ε4 carriers was estimated, and 90% confidence intervals were computed. Statistical equivalence was established when the confidence interval fell entirely within the predefined bounds, and both one-sided tests were significant at α = .05.

To further quantify the evidence against APOE genotype effects, we used Bayesian hypothesis testing [68]. For each outcome measure, a null model assuming no difference between APOE genotype groups was compared to an alternative model allowing for a difference in performance between groups, expressed in standard deviation units. Bayes factors were computed for two-sided tests using a Cauchy prior, the default medium prior implemented in the ttestBF function of the BayesFactor package, centred at zero with a scale parameter of r = 0.707. Bayes factors in favour of the null hypothesis (BF_01_) are reported.

### Bootstrapping

To estimate mean differences among other possible APOE genotype groups (ε3ε3, ε3ε4, ε4ε4, ε2ε2, ε2ε3, ε2ε4), we used a non-parametric bootstrapping approach to accommodate unequal group sizes, including small samples for less frequent genotypes. For each outcome measure (including the main SHQ wayfinding outcomes and measures of the other cognitive tasks), pairwise contrasts between APOE ε3ε3 carriers (our reference group) and the remaining genotype groups were computed by resampling participants within each group with replacement. For each contrast, 10,000 bootstrap resamples were drawn, and the mean difference between groups was calculated for each resample. The resulting bootstrap distributions were used to derive percentile-based 95% confidence intervals for the mean differences, defined by the 2.5th and 97.5th percentiles of the bootstrap distribution. A genotype contrast was considered to provide evidence of a group difference when the corresponding confidence interval did not include zero. This analysis was exploratory, and no correction for multiple comparisons was applied.

## Data availability

Source data to reproduce figures and tables will be openly available at the Open Science Framework once the manuscript is accepted for publication (https://osf.io/9vs3f).

## Code availability

All analysis is based on openly available software or custom code, which can be accessed at the Open Science Framework upon acceptance of the manuscript for publication (https://osf.io/9vs3f).

## Acknowledgements

This research was funded in part by the Austrian Science Fund (FWF) [10.55776/P34775], and by the European Research Council (Starting Grant 101164099), awarded to I.C.W. For the purpose of open access, the author has applied a CC BY public copyright license to any author accepted manuscript version arising from this submission.

## Author contributions

Conceptualization: L.P.G. and I.C.W.; Methodology: L.P.G, L.S., C.G., and I.C.W.; Software: L.P.G, L.S., and I.C.W.; Validation: L.P.G.; Formal Analysis: L.P.G., L.S., C.G.; Investigation: L.P.G.; Resources: I.C.W.; Data Curation: L.P.G. and L.S.; Writing – Original Draft: L.P.G. and I.C.W.; Writing – Reviewing & Editing: all authors; Visualization: L.P.G.; Supervision: I.C.W.; Project Administration: I.C.W.; Funding Acquisition: I.C.W.

## Supplementary Results

### S1: Outlier exclusion does not change results for wayfinding distance and duration

To assess the robustness of the two primary wayfinding outcomes (wayfinding distance and duration), we recomputed group comparisons after excluding extreme outliers (± 3 × median absolute deviation, MAD). Results remained virtually identical to the main analysis (wayfinding distance: n_ε3ε3_ = 308 [19 excluded], n_ε3ε4_ = 87 [1 excluded], wayfinding duration: n_ε3ε3_ = 313 [14 excluded], n_ε3ε4_ = 88 [0 excluded], all p > .05).

### S2: Joint wayfinding performance confirms no APOE genotype effect on goal-directed navigation

Although wayfinding distance and duration are closely related measures (i.e., individuals who take shorter paths are also generally faster), they may reflect partially distinct aspects of navigation performance in individual cases. For example, participants may take relatively short routes while requiring more time to navigate, or move quickly while following less direct paths. To capture goal-directed spatial navigation ability in a single composite score accounting for such cases, we created a joint measure combining both variables (see Methods), with higher values indicating better performance (i.e., shorter, faster routes). In line with the substantial correlation between wayfinding distance and duration (Pearson’s r = 0.87, p < .0001; Figure 2C), joint wayfinding performance did not significantly differ between APOE genotype groups (n_ε3ε3_ = 327, n_ε3ε4_ = 88; Wilcoxon rank-sum test, W = 15048, p = .75, r = 0.032; Supplementary Fig. S1). Overall, these results suggest that goal-directed spatial navigation in healthy young adults was not affected by APOE genotype.

### S3: APOE genotype does not affect SHQ performance differently in biologically female and male participants

Biological sex showed a clear main effect on goal-directed spatial navigation performance, whereas APOE genotype did not. This is consistent with previous findings of biological sex differences in wayfinding duration, with men completing wayfinding levels faster than women [1]. Moreover, this raises the question of whether a genotype-related effect might differ across biological sexes, which would not be captured by a model estimating main effects alone. To test this possibility, and in line with our preregistered analysis plan, we fitted linear regression models that included an APOE × biological sex interaction term. Across the three main wayfinding outcomes (wayfinding distance, wayfinding duration, and mean distance-to-border), there was no evidence for an APOE × biological sex interaction (all p > .05). In other words, the association between APOE genotype and the three main wayfinding outcomes did not differ between biological males and females. We next examined whether including demographic and cognitive covariates altered this pattern by fitting covariate-adjusted interaction models that included biological sex, age, gaming experience, self-reported navigation ability, and map view duration. The APOE × biological sex interaction remained non-significant across wayfinding outcomes (all p > .05), and covariate adjustment did not change the magnitude or direction of the interaction term. These results indicate that APOE genotype does not differentially affect goal-directed spatial navigation performance in biological males and females, irrespective of demographic, cognitive, or task-related factors.

### S4: Equivalence testing and Bayes factors corroborate null effects in the SHQ game

Across all SHQ outcomes, we did not observe significant performance differences between the two APOE genotype groups. However, the absence of a significant effect is difficult to interpret, as a null result may reflect either a true absence of a group effect (meaning that APOE genotype did not affect spatial navigation in healthy young adults) or a lack of statistical power (meaning that any effect of APOE genotype was too small to be detected in our data [2]). We, therefore, next tested whether the observed null effects reflected statistical equivalence between APOE genotype groups. To address this within the frequentist framework, we adopted an equivalence testing approach, which allows researchers to evaluate whether an effect is small enough to be considered statistically equivalent [3]. To do so, we conducted two one-sided tests (TOST) for each outcome, using standardised equivalence bounds of ± 0.3 Cohen’s d [2,4]. If the two genotype groups were truly equivalent in their performance, the confidence intervals for the group differences would fall entirely within the specified bounds. For wayfinding distance, wayfinding duration, and mean distance-to-border, the 90% confidence intervals were fully contained within these bounds, and both one-sided tests were significant (all p < .05; Figure 3C). In other words, and in line with our results so far, the two genotype groups performed within a statistically equivalent range.

To complement our analysis, we next used Bayes factors to quantify the strength of evidence for the absence of APOE genotype performance differences. While the frequentist analyses showed no significant group differences and equivalence testing indicated that the null results reflected statistical equivalence, Bayes factors allow a direct evaluation of whether the data provide stronger support for the null model or for the alternative [5]. We used the default medium prior from the ttestBF function of the BayesFactor package, which is a Cauchy prior on the standardised effect size, centred at zero with a width of r = 0.707. This distribution favours small over large effect sizes [5], consistent with our expectation that potential effects of APOE genotype on spatial navigation in healthy young adults may be subtle [6]. Across all three main wayfinding outcomes, Bayes factors supported the null model over the alternative, with BF_01_ values indicating moderate evidence for the absence of APOE genotype differences (wayfinding distance: 6.34, wayfinding duration: 7.58, and mean distance to border: 6.88, Figure 3C). Altogether, these results suggest that goal-directed spatial navigation ability in healthy young adults is unaffected by the APOE ε4ε3 genotype.

### S5: Equivalence testing and Bayes factors corroborate null effects in tasks assessing working memory, processing speed, executive function, and face recognition

Across all cognitive tasks assessing working memory, processing speed, executive function, and face recognition, the two APOE genotype groups did not differ significantly in their performance. To determine whether these null effects reflected a true absence of group differences rather than insufficient statistical power to detect small effects, we conducted equivalence tests. Following established procedures, we applied two one-sided tests (TOST) for each task measure, using standardised equivalence bounds of ± 0.3 Cohen’s d [2,3]. For SSTM, TMT B, and CFMT, the 90% confidence intervals of the mean differences fell entirely within the specified bounds, and both one-sided tests were significant (all p < .05; Figure 4C). This suggests that any group differences in these tasks were smaller than the predefined equivalence range and can therefore be considered statistically equivalent. Scores of the MU task and the TMT A slightly exceeded the equivalence bounds, allowing for two different interpretations. Either APOE genotype groups performed similarly on these tasks, though not statistically equivalent by our criteria, or any true performance differences were minimal (and thus too small for us to detect with sufficient power). Together, these findings support the notion of equivalent performance on SSTM, TMT B, and CFMT across APOE genotype groups, while the evidence for MU and TMT A is less conclusive. We speculate that if any differences in MU and TMT A performance truly exist, they are likely very small.

We next turned to Bayesian statistics to examine how strongly the data supported the absence of APOE genotype differences in performance across the cognitive tasks. Specifically, we compared a null model assuming no difference between APOE genotype groups to an alternative model estimating a possible difference in standardised effect size. As above, we used the default medium prior implemented in the ttestBF function of the BayesFactor package, corresponding to a Cauchy prior on the standardised effect size centred at zero with a width of r = 0.707. We calculated Bayes factors for each task. Across all task measures, the resulting BF_01_ values indicated a preference for the null model over the alternative (that is, the data were more likely under the assumption of no group differences), providing moderate evidence for the absence of APOE-related performance differences (MU: 4.72, SSTM: 7.37, TMT A: 3.14, TMT B: 6.39, CFMT: 6.32; Figure 4C). Overall, these results indicate that the APOE genotype did not affect participants’ working memory, executive function, or face recognition.

## Supplementary Methods

### S1: Preregistration and deviations

The study was preregistered on OSF on July 11, 2023 (https://doi.org/10.17605/OSF.IO/ZMDBF). We deviated from the preregistered analysis plan in six aspects.

First, the preregistration specified that APOE ε3ε4 and ε4ε4 carriers would be analysed as a single APOE ε4 carrier group and compared with ε3ε3 carriers, to test for potential ε4-related effects on cognition in a healthy young cohort. In the analyses reported here, we opted for keeping these genotype groups separate. Although both groups included at least one ε4 allele, ε3ε4 and ε4ε4 carriers are known to differ in their associated Alzheimer’s disease risk, with ε4ε4 carriers showing markedly higher risk than ε3ε4 carriers [7]. Treating these genotypes as a single group could therefore obscure dose-dependent or genotype-specific effects in two distinct ways: an effect driven mainly by ε4ε4 carriers could appear diluted (and statistically weaker) when pooled with ε3ε4 carriers, or, conversely, an effect specific to ε4ε4 carriers could be misattributed to the ε4 carrier group as a whole, falsely suggesting generalisation to ε3ε4 carriers. We therefore focused the main analyses on the comparison between ε3ε4 and ε3ε3 carriers, in line with previous studies investigating the effect of APOE ε4 on cognition [8,9]. As a sensitivity check, we repeated the analysis using the preregistered ε4 carrier grouping. This yielded virtually unchanged results, with no significant performance differences between ε3ε3 carriers and the combined ε4 carrier group (n_ε3ε3_ = 327, n_ε4_ = 99; wayfinding distance: p = .17, wayfinding duration: p = .29, mean distance-to-border: p = .28).

Second, we used linear regression models rather than the preregistered linear mixed-effects models for the follow-up analyses of the APOE × biological sex interaction. Linear mixed-effects models were originally planned to account for repeated observations within participants across SHQ levels. Here, we tested the APOE × biological sex interaction on outcomes aggregated across levels, following directly from our primary genotype comparisons (Wilcoxon rank-sum tests), which were likewise conducted on aggregated data. The same approach was used for the covariate-adjusted interaction models, which included biological sex, age, gaming experience, self-reported navigation ability, and map viewing duration. To separately assess whether any APOE-related differences were tied to a specific SHQ level, which is a question that the aggregated models could not address, we additionally fitted linear mixed-effects models. Game level, APOE genotype, and their interaction were treated as fixed effects, for each of the three main wayfinding outcomes (wayfinding distance, wayfinding duration, and mean distance-to-border), as originally planned. Across all three outcomes, we observed no significant APOE genotype × game level interaction (all p > .05), suggesting that APOE-related differences did not vary by game level.

Third, we had planned to examine the interaction between APOE genotype and biological sex for wayfinding distance, wayfinding duration, and flare accuracy. During flare accuracy levels, participants had to indicate their starting position after navigating to a target location, allowing us to assess path integration ability. Since prior work had linked path integration to APOE ε4-related differences in spatial navigation, we expected ε4 carriers to show early behavioural effects on flare accuracy specifically [8,9]. However, flare accuracy scores were discrete and strongly skewed toward the maximum, with 80% of participants performing at ceiling. We therefore retained flare accuracy in the primary group comparison but did not carry it forward into the follow-up regression models, as the restricted range left too little interindividual variability to meaningfully test effects of APOE genotype, biological sex, or their interaction.

Fourth, we deviated from our preregistered plan when deciding which demographic, cognitive, and questionnaire-based variables to include in the follow-up models. The preregistered procedure was exploratory and would have involved adding and removing predictors based on model fit in a stepwise fashion. In the final analyses, we instead decided to use a fixed set of covariates selected for their theoretical or task-related relevance to navigation performance. Biological sex and age have been consistently linked to individual differences in spatial navigation [10–12], alongside self-reported navigation ability (Santa Barbara Sense Of Direction Scale, SBSOD [13]) and computer- and smartphone-based gaming experience involving virtual navigation. We also included map viewing duration, as longer inspection times may have helped participants memorise checkpoint locations. This fixed-covariate approach allowed us to test whether the APOE ε4 result changed after accounting for these variables. The results were consistent with the primary analyses: APOE ε4 status was not associated with any of the three main wayfinding outcomes (wayfinding distance, wayfinding duration, mean distance-to-border).

Fifth, we used Wilcoxon rank-sum tests instead of the preregistered independent-samples t-tests for the main group comparisons of wayfinding distance, wayfinding duration, and mean distance-to-border. Wayfinding distance and wayfinding duration significantly deviated from the normal distribution (Shapiro-Wilk tests, both p < .05, consistent across APOE subgroups) and were therefore analysed nonparametrically. Mean distance-to-border met the normality assumption, both in the pooled sample and within each APOE subgroup, but we applied the same nonparametric test for comparability and consistency across the three main wayfinding outcomes. Given our directional hypotheses that ε3ε4 carriers would show poorer navigation performance, reflected in longer wayfinding distances and durations as well as navigation at closer distances to borders of the virtual environment, we applied one-tailed tests to these outcomes. Nevertheless, no group differences emerged. Notably, using t-tests did not change the results (n_ε3ε3_ = 327, n_ε4_ = 88; wayfinding distance: p = .27, wayfinding duration: p = .49, mean distance-to-border: p = .33).

Sixth, for the main group comparison of flare accuracy specifically, we conducted Fisher’s exact test rather than the Wilcoxon rank-sum test. Flare accuracy scores were calculated as the sum of three per-level scores (1-3 points each, range 3-9) and were strongly skewed toward the maximum (s. deviation 3). This left little variability for a rank-based test to discriminate between groups, so we instead chose Fisher’s exact test, which is well-suited to a small number of discrete score categories.

### S2: Model assumptions

Model assumptions for the linear regression and linear mixed-effects models reported in the main text were assessed using graphical diagnostics. For each model, we inspected residual Q-Q plots to assess residual normality and plotted residuals against fitted values to evaluate homoscedasticity and linearity. We report diagnostics separately for four models: the linear regression model testing effects of APOE genotype and covariates on SHQ performance, the linear regression model testing the APOE genotype × biological sex interaction, the covariate-adjusted linear regression model testing the APOE genotype × biological sex interaction, and the linear mixed-effects model testing the APOE genotype × game level interaction. For each model, diagnostics are shown separately for the three main wayfinding outcomes (wayfinding distance, wayfinding duration, and mean distance-to-border; Supplementary Fig. 2-5).

We first examined residual normality using Q-Q plots. Across the three regression models (Supplementary Fig. 2-4), wayfinding distance and wayfinding duration deviated from normality, with a heavier upper tail than expected, consistent with the non-normal distribution of these two outcomes (Shapiro-Wilk test; see Supplementary Methods S1). Mean distance-to-border did not deviate from normality (Shapiro-Wilk test; see Supplementary Methods S1). In the linear mixed-effects model (Supplementary Fig. 5), the upper-tail deviation from normality was more pronounced for wayfinding distance and duration than in the other three models, potentially reflecting the use of level-wise rather than aggregated data. Mean distance-to-border, which had not deviated from normality in the regression models, showed a mild deviation at both tails in this model.

We next examined plots of residuals against fitted values to assess homoscedasticity and linearity. Plots did not raise homoscedasticity concerns for mean distance-to-border, whereas for wayfinding distance and duration, residual spread increased slightly at higher fitted values across models. In the linear regression model testing the APOE genotype × biological sex interaction (Supplementary Fig. 3), which included only categorical predictors, fitted values formed discrete clusters rather than a continuous range. Mean distance-to-border in the linear mixed-effects model (Supplementary Fig. 5) showed a similar pattern, with fitted values clustering by game level. The residual variance for this outcome was comparatively low, suggesting that game level may have been the main driver of the fitted values. Residual spread was comparable across clusters in the biological sex interaction model, but varied between game-level clusters in the mixed-effects model.

Diagnostics for the covariate model (Supplementary Fig. 2) and the covariate-adjusted interaction model (Supplementary Fig. 4) were nearly identical across outcomes, since the two models differed only in the small, non-significant APOE × biological sex interaction term. Accordingly, fitted values from the two models differed only marginally (e.g., for wayfinding distance: mean relative difference between the two models’ fitted values < 1%). Overall, the diagnostics revealed some deviations, mainly for wayfinding distance and duration. These outcomes showed a heavier upper tail than expected and a slight increase in residual spread at higher fitted values across all four models, both most pronounced in the mixed-effects model. Mean distance-to-border met assumptions closely in the three regression models, departing only in the clustering of fitted values by game level in the mixed-effects model.

## Supplementary Figures

**Fig. S1:**
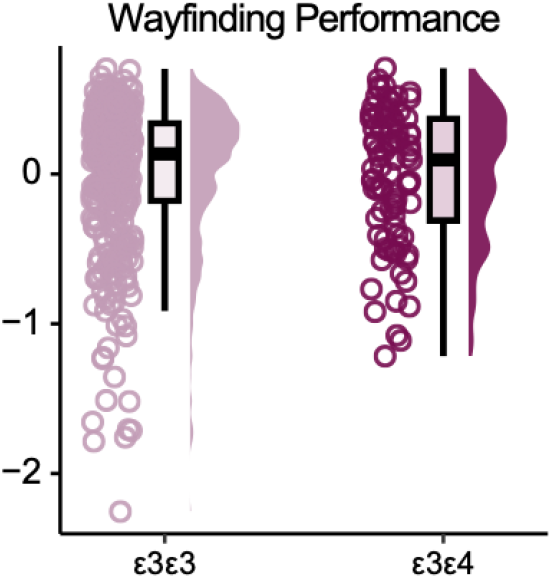
No difference in joint wayfinding performance between genotype groups. Data points show individual joint wayfinding performance scores (combined z-standardised wayfinding distance and duration, multiplied by - 0.5, such that higher values indicate better performance) for APOE ε3ε3 carriers (in rose) and ε3ε4 carriers (in dark red). Boxplots mark the median, the interquartile range, and whiskers indicate the minimum and maximum outlier values. Group differences were assessed using a Wilcoxon rank-sum test (p = .75).

**Fig. S2:**
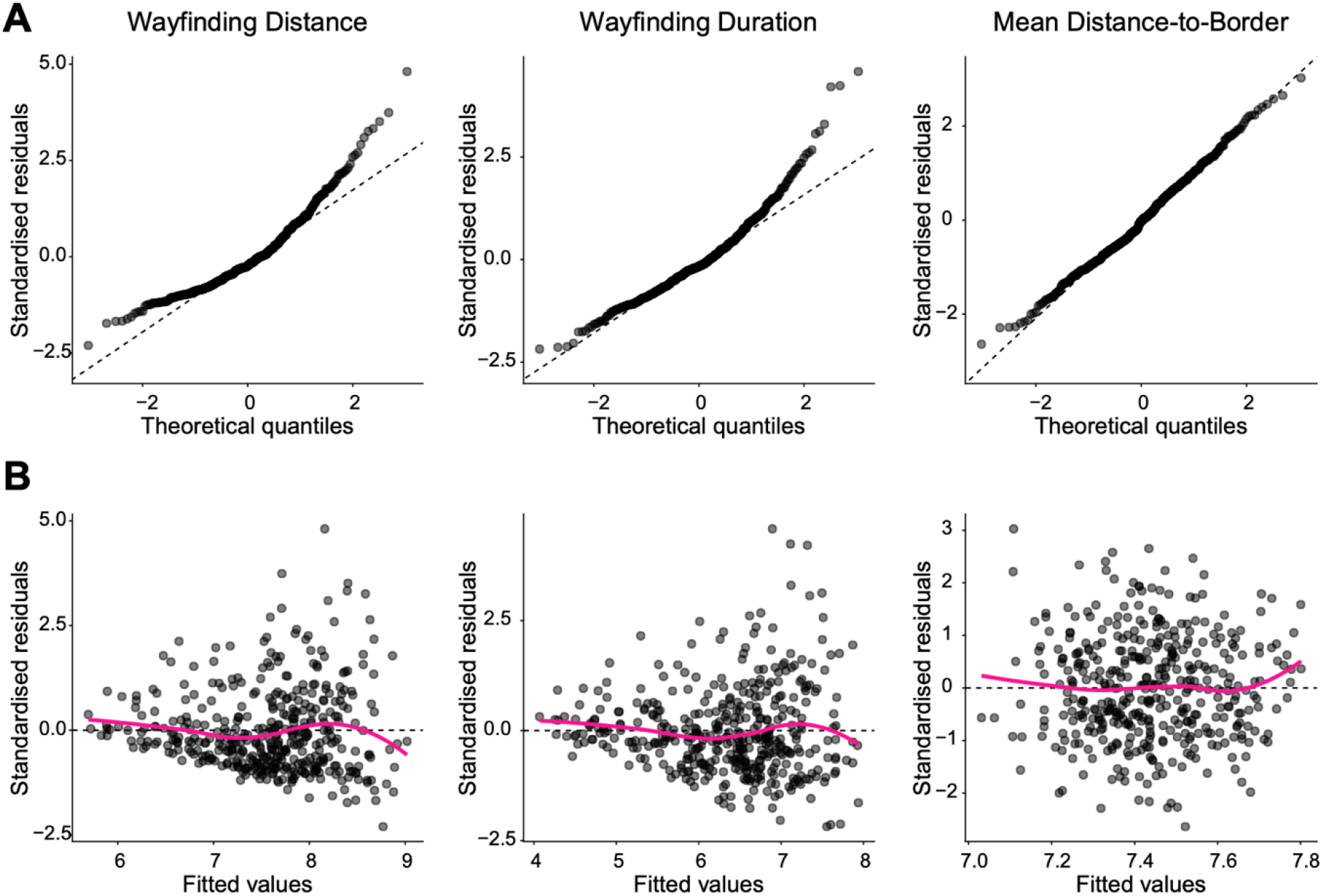
Model assumptions for the linear regression model testing effects of APOE genotype and covariates on SHQ performance. A. Residual Q-Q plots for wayfinding distance (left), wayfinding duration (middle), and mean distance-to-border (right), from linear regression models including APOE genotype, biological sex, age, gaming experience, self-reported navigation ability, and map viewing duration as fixed effects. B. Standardised residuals plotted against fitted values for wayfinding distance (left), wayfinding duration (middle), and mean distance-to-border (right). Dashed lines indicate the Q-Q reference line and the zero line for residuals; solid lines show a smoothed trend through the residuals.

**Fig. S3:**
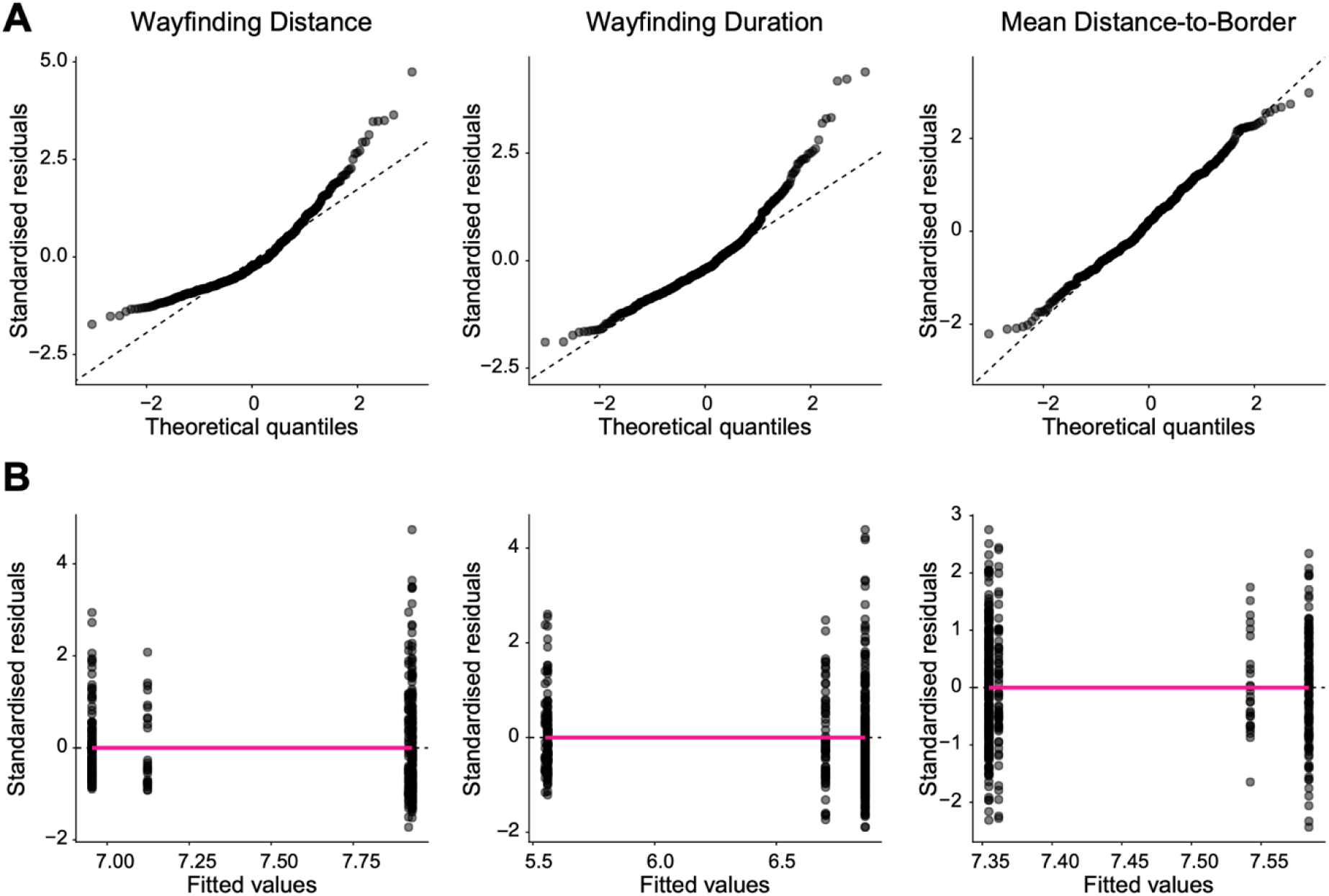
Model assumptions for the linear regression model testing the APOE genotype × biological sex interaction on SHQ performance. A. Residual Q-Q plots for wayfinding distance (left), wayfinding duration (middle), and mean distance-to-border (right). B. Standardised residuals plotted against fitted values for wayfinding distance (left), wayfinding duration (middle), and mean distance-to-border (right). Dashed lines indicate the Q-Q reference line and the zero line for residuals; solid lines show a smoothed trend through the residuals.

**Fig. S4:**
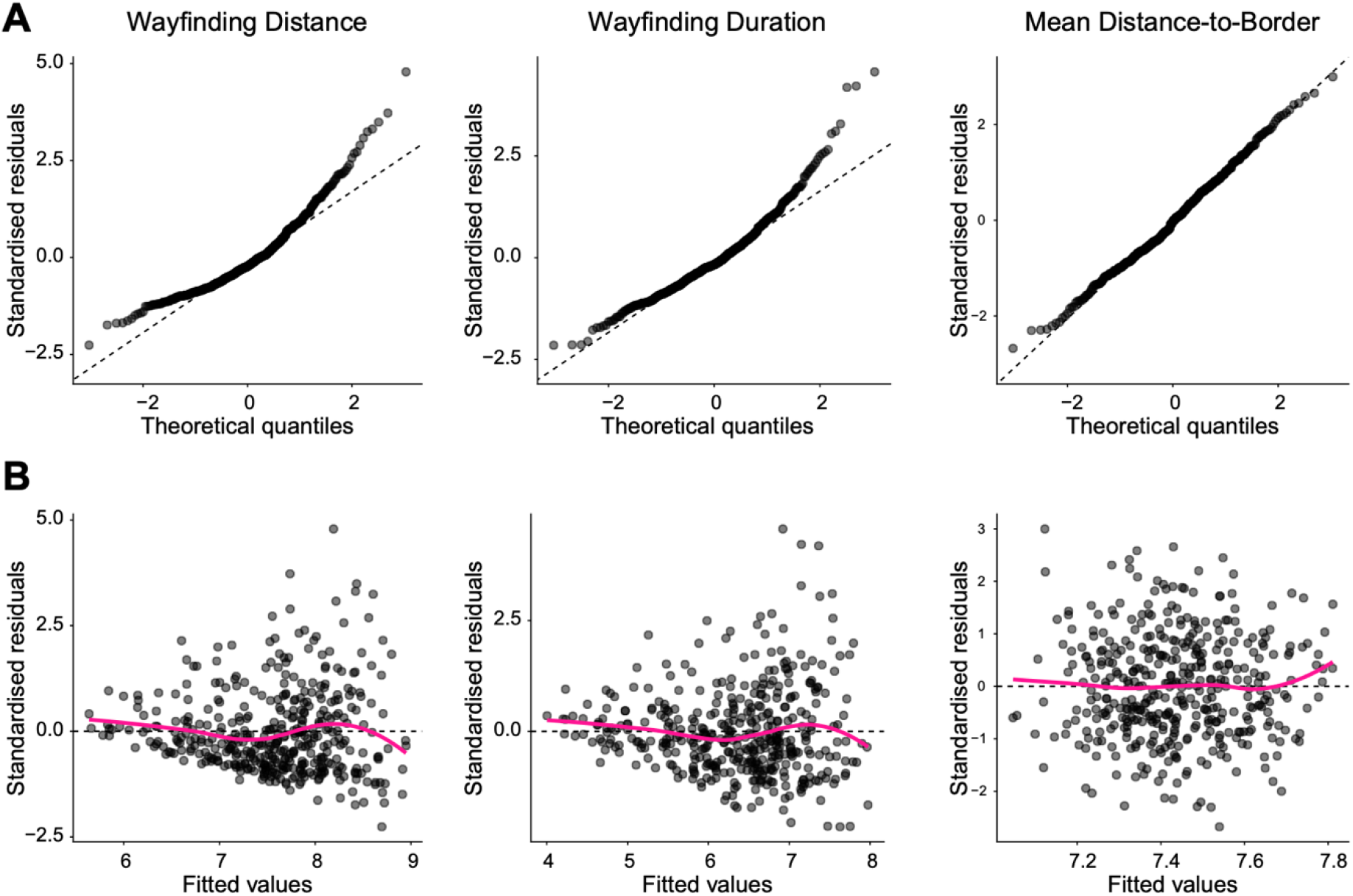
Model assumptions for the covariate-adjusted linear regression model testing the APOE genotype x biological sex interaction on SHQ performance. A. Residual Q-Q plots for wayfinding distance (left), wayfinding duration (middle), and mean distance-to-border (right), from models including biological sex, age, gaming experience, self-reported navigation ability, and map viewing duration as covariates. B. Standardised residuals plotted against fitted values for wayfinding distance (left), wayfinding duration (middle), and mean distance-to-border (right). Dashed lines indicate the Q-Q reference line and the zero line for residuals; solid lines show a smoothed trend through the residuals.

**Fig. S5:**
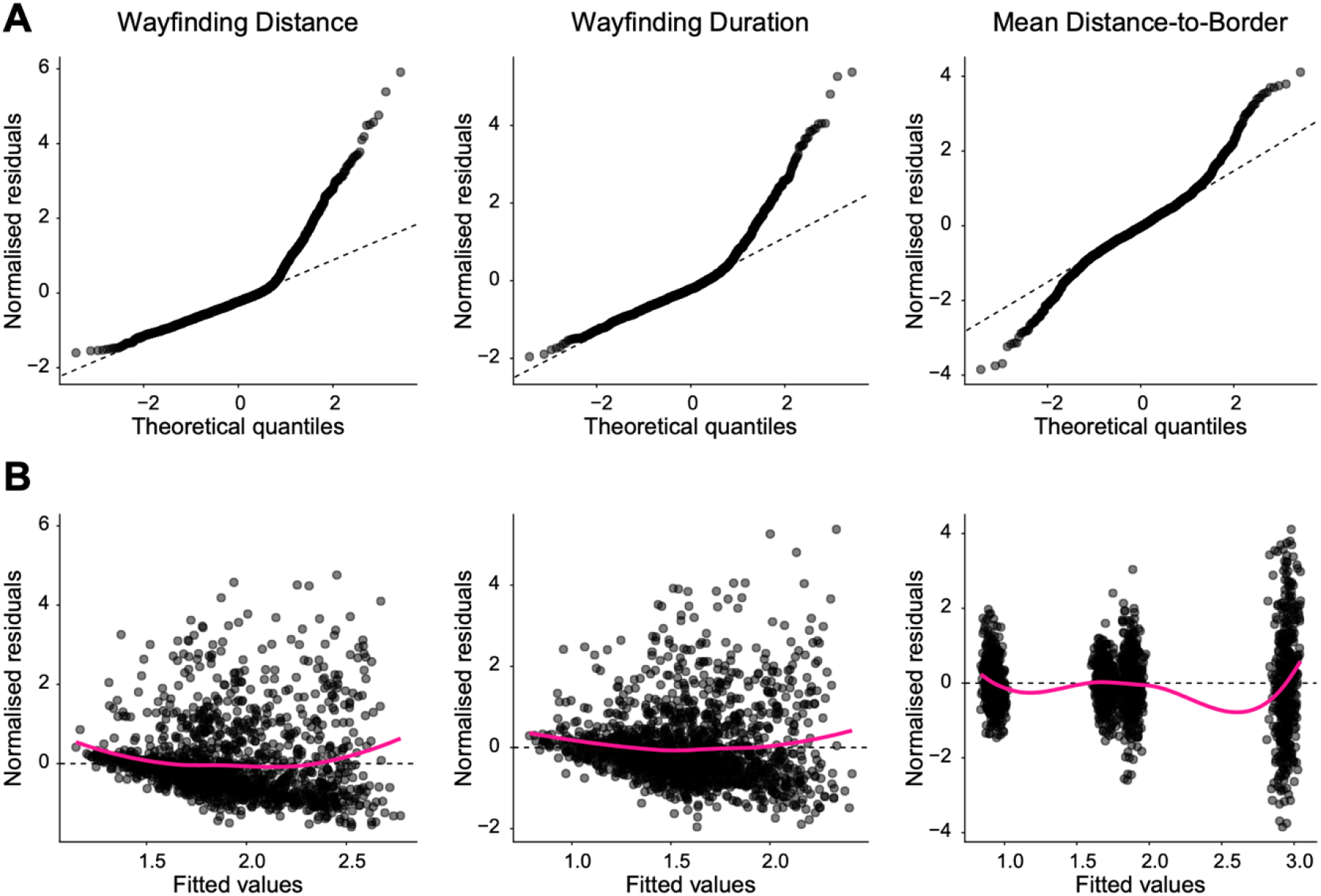
Model assumptions for the linear mixed-effects model testing the APOE genotype × game level interaction on SHQ performance. A. Residual Q-Q plots for wayfinding distance (left), wayfinding duration (middle), and mean distance-to-border (right), from models including game level, APOE genotype, and their interaction as fixed effects, together with biological sex, age, gaming experience, self-reported navigation ability, and map viewing duration as covariates, and random intercepts for participants. B. Normalised residuals plotted against fitted values for wayfinding distance (left), wayfinding duration (middle), and mean distance-to-border (right). Dashed lines indicate the Q-Q reference line and the zero line for residuals; solid lines show a smoothed trend through the residuals.

## Supplementary Tables

**Table 1.**
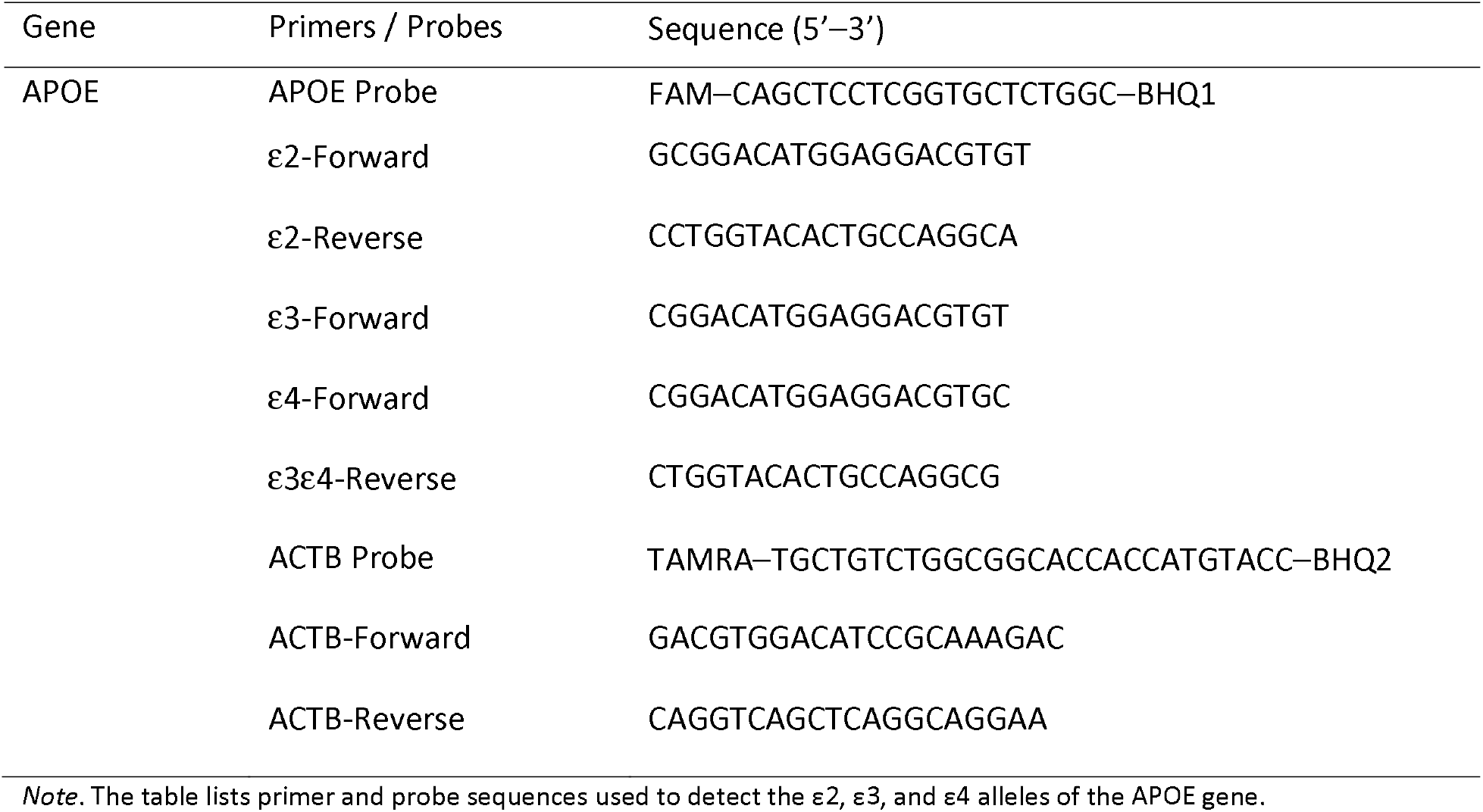
Sequences of primers and probes for APOE genotyping.

